# Intrinsically disordered RNA-binding motifs cooperate to catalyze RNA folding and drive phase separation

**DOI:** 10.1101/2024.03.27.586925

**Authors:** Annika Niedner-Boblenz, Thomas Monecke, Janosch Hennig, Melina Klostermann, Mario Hofweber, Elena Davydova, André P. Gerber, Irina Anosova, Wieland Mayer, Marisa Müller, Roland Gerhard Heym, Robert Janowski, Jean-Christophe Paillart, Dorothee Dormann, Kathi Zarnack, Michael Sattler, Dierk Niessing

## Abstract

RNA-binding proteins are essential for gene regulation and the spatial organization of cells. Here, we report that the yeast ribosome biogenesis factor Loc1p is an intrinsically disordered RNA-binding protein with eight repeating <u>p</u>ositively charged, <u>u</u>nstructured <u>n</u>ucleic acid binding (PUN) motifs. While a single of these previously undefined motifs stabilizes folded RNAs, multiple copies strongly cooperate to catalyze RNA folding. In the presence of RNA, these multivalent PUN motifs drive phase separation. Proteome-wide searches in pro-and eukaryotes for proteins with similar arrays of PUN motifs reveal a strong enrichment in RNA-mediated processes and DNA remodeling. Thus, PUN motifs are potentially involved in a large variety of RNA-and DNA-related processes by concentrating them in membrane-less organelles. The general function and wide distribution of PUN motifs across species suggests that in an ancient “RNA world” PUN-like motifs may have supported the correct folding of early ribozymes.

## Introduction

Biogenesis of eukaryotic ribosomes requires the correct processing and assembly of four ribosomal RNAs (rRNAs; 5S, 5.8S, 25S, and 18S) with 79 ribosomal proteins. In the budding yeast *Saccharomyces cerevisiae*, this process is guided by 76 snoRNAs and more than 200 different assembly factors (1). Ribosomal proteins stabilize folded rRNAs, function as chaperones to favor the correct rRNA fold, or alter the rRNA fold to create binding sites for other ribosomal proteins (2). In contrast to ribosomal proteins, so-called assembly factors are non-integral, associated components of the ribosome (1). They often share properties with ribosomal proteins such as unusually high isoelectric points (**Table S1**). However, assembly factors are functionally much more diverse, for instance including endo-and exonucleases, protein-and RNA-modifying enzymes, RNA helicases and other ATPases, GTPases, kinases and phosphatases, RNA-binding proteins, and putative scaffolding proteins (3). Often their individual molecular function has been assigned to a particular step of RNA processing, export, or maturation, based on the observed depletion phenotypes.

In *S. cerevisiae*, the nucleolar RNA-binding protein Loc1p is required for efficient ribosome biogenesis (4). It also has been suggested to be necessary for the generation of distinct classes of ribosomes with different sets of ribosomal protein paralogs (5). In addition, Loc1p fulfills a second function. It supports *ASH1* mRNA localization to the bud tip by stabilizing the formation of immature messenger ribonucleoprotein particles (mRNPs) in the nucleus (6–8).

Loc1p is a constituent of pre-60S ribosomal particles (9–12) and co-purifies with Nop7, which functions in 90S and 66S ribosome biogenesis (4) and with the ribosomal protein Rpl43 (13,14). Consistently, a *loc1Δ* strain showed reduced 60S subunit levels and half-mer polyribosomes in sucrose gradients (15). A closer examination of rRNA processing in the *loc1Δ* strain revealed impaired cleavage at sites A_0,_ A_1_ and A_2_ (15). Nevertheless, a functional understanding of the molecular role of Loc1p in the biogenesis of ribosomes is still missing.

Here, we report that Loc1p is an intrinsically disordered protein (IDP) with strong, ATP-independent RNA-annealing activity. We identified eight short, <u>p</u>ositively charged, <u>u</u>nfolded <u>n</u>ucleic acid-binding (PUN) motifs in Loc1p, which cooperatively act as potent RNA annealers. Furthermore, multiple PUN motifs mediate RNA-driven phase separation, suggesting a role of PUN motif-containing proteins (PUN proteins) in the subcellular organization of RNAs. Several nuclear proteins involved in RNA-metabolic processes and neurogenesis in higher eukaryotes are highly enriched with multiple PUN motifs. Together, these findings imply that PUN proteins are core constituents at the interface of protein-nucleic acid complexes across species. We speculate that in ancient times such short repetitive elements with PUN-like properties could have fostered the transition from purely RNA-based macromolecules to an RNP-dominated world.

## Material and Methods

### Protein expression and purification

MBP-tagged and His_6_-tagged Loc1p (UniProt ID P43586) was expressed and purified as previously described (6). Loc1p (1-20)_10_ was expressed with an N-terminal His_6_ tag in *E. coli* strain BL21 (DE3) Star. Expression and purification of Loc1p (1-20)_10_ was identical to the procedures of wild-type Loc1p (6). Loc1p (3-15) was expressed with an N-terminal GST tag in *E. coli* strain BL21-Gold (DE3) pRARE using M9 minimal media supplemented with ^15^N ammonium chloride and ^13^C glucose. After induction with 0.25 mM isopropyl β-D-1-thiogalactopyranoside (IPTG) in the mid-logarithmic growth phase the cells were cultured for 16 h at 18 °C and subsequently harvested. Cells were resuspended in lysis buffer (20 mM Na_2_HPO_4_/NaH_2_PO_4_ pH 7.4; 500 mM NaCl; EDTA-free complete protease inhibitor (Roche); 0.4 mM PMSF) and sonicated at 4 °C. After centrifugation the clarified supernatant was applied to a GSTrap FF sepharose column (GE Healthcare) and washed with high-salt (1 M NaCl) and low-salt buffer (150 mM NaCl). The protein was digested with thrombin over night at 4 °C and further purified by cation-exchange chromatography. Peptide-containing fractions were pooled, dialyzed in water, and lyophilized.

*Yersenia enterocolitica* Hfq (UniProt ID Q56928) was expressed N-terminally His_6_-SUMO-tagged in *E. coli* strain ER 2566. At mid-logarithmic growth phase, the expression was induced by addition of 0.25 mM IPTG and cells were cultured for 16 h at 18 °C and harvested. Cells were sonicated at 4 °C in lysis-buffer (100 mM HEPES pH 8; 200 mM MgCl_2_; 1 M NaCl; 1 mM DTT; EDTA-free complete protease inhibitor (Roche)). After centrifugation, the supernatant was applied to a HisTrap-FF sepharose column (GE Healthcare) and washed with low-salt buffer (500 mM NaCl) before the protein was eluted with 500 mM imidazole in low-salt buffer. The protein was further purified using size exclusion chromatography (Superdex S75; GE Healthcare).

Imp4p (UniProt ID P53941) was expressed His_6_-SUMO-tagged in *E. coli* strain BL21 (DE3) Star. When cells reached the mid-logarithmic growth phase, expression was induced by addition of 0.25 mM IPTG. After 16 h of culture at 18 °C the cells were harvested and resuspended in lysis buffer (20 mM HEPES pH 7.8; 500 mM NaCl; 1 mM DTT) before sonication was performed at 4 °C. The suspension was centrifuged and the clarified supernatant applied to HisTrap-FF sepharose column. After washing steps with high-salt (1 M NaCl) and low-salt buffer (200 mM NaCl) the protein was eluted with 500 mM imidazole and further purified using size-exclusion chromatography (Superose S12 column; GE Healthcare).

Coding sequences of Loc1p (1-20)_3_, Loc1p (1-20)_5_ and SRRM1 (amino acids 151-377) (UniProt ID Q8IYB3) were cloned into pGEX4T1 vector with an N-terminal GST-tag followed by a thrombin cleavage site. Loc1p (1-20)_3_, Loc1p (1-20)_5_ and SRRM1 were expressed in *E. coli* BL21-Gold (DE3) pRARE using auto inducing medium (16). The cells were grown for 4 h at 37 °C, 18h at 20 °C and then harvested at 10,000 x g and 4 °C for 30 min. For each protein purification protocol was identical. The pellet from 2 l of culture was resuspended in buffer A (20 mM Tris pH 8.5, 500 mM NaCl), supplemented with DNase I and EDTA-free protease inhibitor cocktail (Roche) to a total volume of 50 ml. After sonication, the lysate was clarified by centrifugation (20,000 x g, for 30 min), loaded onto 5 ml GST-trap column (GE Healthcare), equilibrated in buffer A, and washed with 20 column volumes of buffer A. High salt wash was implemented (10 column volumes; 20 mM Tris pH 8.5, 1 M NaCl). GST-tagged protein was eluted from the column in 10 CV linear glutathione gradient using buffer B (20 mM Tris pH 8.5, 500 mM NaCl, 10 mM glutathione). Fractions with protein were combined and dialyzed against 20 mM Tris pH 8.5, 150 mM NaCl, 2.5 mM CaCl_2_ o/n at 4 °C with Thrombin added to remove the GST-tag (1:100 ratio). Thrombin, uncleaved protein as well as cleaved GST were removed by applying everything onto a 5 ml SP column (GE Healthcare) equilibrated with 20 mM Tris pH 8.0, 50 mM NaCl buffer. Prior to loading the sample onto SP column, sample buffer was exchanged to 20 mM Tris pH 8.0, 50 mM NaCl to decrease salt concentration. The protein of interest was eluted from the column in 10 CV linear salt gradient using 20 mM Tris pH 8.0, 1 M NaCl. The peak fractions were analyzed using 12% SDS-PAGE. The fractions of interest were combined, concentrated to 5 ml, and applied on a Superdex200 16/60 column (GE Healthcare) equilibrated with 20 mM HEPES pH 7.5, 200 mM NaCl, 2 mM MgCl_2_. The selected fraction from the size exclusion step were then pooled and concentrated by ultrafiltration to 4.7 mg/ml (Loc1p (1-20)_3_), 1.8 mg/ml (Loc1p (1-20)_5_), and 1.5 mg/ml (SRRM1).

### Labeling of His_6_-Loc1p-S7C by Maleimide-Cy5

The single cysteine mutant His_6_-Loc1p-S7C (Loc1p_S7C_) was purified as described for the wild-type protein (see above) and site-specifically labeled using Cy5 Maleimide mono-reactive dye (Cytiva). Prior to labeling, 1 mg purified Loc1p_S7C_ was desalted (150 mM NaCl, 10 mM HEPES pH 7.2), flushed with gaseous N_2_, and incubated for 30 min at 20 °C. A 100-fold molar excess of TCEP was added, flushed with N_2_, and incubated for another 10 min at 20 °C. Cy5-maleimide was dissolved in 50 µl N,N-dimethylformamide, mixed with the protein solution, and flushed with N_2_ again. The labeling reaction was incubated at 20 °C for 2 h and at 4 °C overnight. The reaction was stopped by the addition of 5 mM DTT and labeled protein was separated from free dye by means of gel filtration (Superdex200 16/60) in 150 mM NaCl, 10 mM HEPES pH 7.2, and 1 mM DTT. Fractions containing labeled protein were pooled, concentrated to 2.2 mg/ml, frozen in liquid nitrogen, and stored at -80 °C.

### Identification of Loc1p-associated RNAs with DNA microarrays

C-terminally TAP-tagged Loc1p was purified from 1 L of cells grown in YPD medium as previously described (17,18). cDNA was synthesized from 3 µg of total RNA derived from the extract and 500 ng of affinity-isolated RNA and labeled with Cy3 and Cy5 fluorescent dyes, respectively. Samples were mixed and hybridized to cDNA microarrays (17,18) and data submitted to the Stanford Microarray Database (SMD) (19). Normalized log_2_ median ratios from three independent Loc1p affinity isolations and four mock control isolations were filtered for regression correlation of >0.6, signal over background >1.8 in the channel measuring total RNA, and only features that met this criterion in >60% of the arrays were considered (total 6,726 features). Data were exported into Microsoft Excel (**Tables S2** and **S3**) for further analysis. Microarray data is available for download via the PUMA database (http://puma.princeton.edu), experiment set No. 7342. SAM analysis was performed as described (18) in R-studio, and overrepresented GO terms among protein coding genes identified with Webgestalt (20).

### RNA and DNA production

All RNAs and DNAs were ordered from IBA, Biomers or Eurofins, respectively. *EAR1* and *ASH1* E3 RNAs were *in vitro* transcribed using the MEGAshortscript T7 transcription kit from life technologies. The sequences of all oligonucleotides used in this study are listed in **Table S4**.

### Electrophoretic mobility shift assay (EMSA)

EMSAs were performed as previously described (6,21). Briefly, in a total volume of 20 µl the indicated protein amount was incubated with 5 nM radioactively labeled RNA and 100 µg/ml competitor yeast tRNA in HNMD buffer (20 mM HEPES, pH 7.8, 200 mM NaCl, 2 mM MgCl_2_, 2 mM DTT) supplemented with 4% glycerol. For competitive EMSA experiments 10-fold, 100-fold, and 500-fold molar excess of the indicated unlabeled RNA was added and yeast tRNA was omitted. Samples were incubated for 25 min at 25 °C before RNA-protein complexes were resolved by native 6% TBE PAGE. Gels were fixed for 15 min in fixing solution (10% (v/v) acetic acid; 30% (v/v) methanol) before vacuum drying. Gels were analyzed using radiograph films. For each EMSA, at least three independent experiments were performed (n ≥ 3).

### Yeast growth assay

YPD pre-cultures of wild-type (MATa, his3ι11, leu2ι10, met15ι10, ura3ι10) and *loc1*ι1 yeast strains (MATa, ade21, can1100, his311,15, leu23,112, trp11, ura3, GAL, psi+, loc1::natNT2) were inoculated from YPAD agar plates and grown for 24 hours at 30 °C and 180 rpm. Main cultures were started at a calculated OD_600_ of 0.05 in 70 ml YPAD media and grown at 30 °C and 180 rpm. Every hour, OD_600_ was measured (undiluted at OD_600_ 0 -0.5 and 1:5 diluted at OD_600_ ≥ 0.5). Data points collected over 10 hours were plotted and fitted to an exponential growth curve for wildtype and *ι1loc1* individually, which was used to calculate doubling times (T_d_ = tau) from three independent experiments.

### Purification of yeast ribosomes

All ribosome-purification steps were performed at 4 °C unless stated otherwise. Wild-type yeast cells (S288c) and *loc1Δ* cells were cultured until reaching an OD_600_ 1.5 to 2.5, washed with water and 1% KCl (w/v), and resuspended in buffer (100 mM Tris pH 8; 10 mM DTT). After 15 min incubation at room temperature the cells were pelleted and resuspended in lysis-buffer (20 mM HEPES pH 7.5; 100 mM KOAc; 7.5 mM Mg(OAc)_2_; 125 mM sucrose; 1 mM DTT; 0.5 mM PMSF; EDTA-free Complete protease inhibitors (Roche)). Cells were disrupted in a microfluidizer (3 times; 20,000 psi) and centrifuged for 15 min at 15,500 rpm (SS34 rotor). The supernatant was again centrifuged for 30 min at 37,000 rpm (Ti-70 rotor) and only the clear fraction (S100) was further purified. Subsequently 15 ml S100 was layered over 4 ml of 2 M sucrose cushion and 4 ml of 1.5 M sucrose cushion (sucrose cushion buffer: 1.5 or 2 M sucrose; 20 mM HEPES pH 7.5; 500 mM KOAc; 5 mM Mg(OAc)_2_; 1 mM DTT; 0.5 mM PMSF) and centrifuged in a Ti 70 rotor tube for 18 h at 53,250 rpm. The ribosome pellet was resuspended in 500 µl water. Experiments were performed at least twice (n ≥ 2).

### Purification of *Mycobacterium smegmatis* ribosomes

All purification steps were performed at 4 °C unless stated otherwise. *Mycobacterium smegmatis* was cultured at 37 °C until an OD_600_ of 0.3-0.8 was reached. The cells were harvested by 15 min centrifugation at 5,000 x g and resuspended in 70S HB buffer (20 mM Tris pH 7.4; 100 mM NH_4_Cl; 10 mM MgCl_2_; 3 mM beta-mercaptoethanol). The cell suspension was supplemented with 2 units DNAse I per gram of cell mass and cells were disrupted using a microfluidizer at 10,000 to 16,000 psi 3 to 5 times. The lysate was cleared two times by centrifugation (30,000 x g, 30 min). NH_4_Cl and beta-mercaptoethanol were added to reach a final concentration of 350 mM and 7 mM, respectively. The lysate was centrifuged again (Ti 70 rotor, 36,600 rpm, 15 min and 15 ml of the clear fraction were loaded onto 9 ml of sucrose cushion (20 mM Tris pH 7.4; 350 mM NH_4_Cl; 10 mM MgCl_2_; 3 mM beta-mercaptoethanol; 1.1 M sucrose). The ribosomes were pelleted using ultracentrifugation (Ti 70 rotor, 36,600 rpm, 17 h), carefully washed three times with 70S buffer (20 mM HEPES pH 7.4; 60 mM NH_4_Cl, 6 mM MgCl_2_; 3 mM beta-mercaptoethanol), and dissolved in 70S buffer by gentle shaking. After short centrifugation, ribosomes were loaded onto 10-40% sucrose gradients in 70S buffer and ultracentrifuged (SW 28 rotor, 16,800 rpm, 16 h). Fractions containing 70S ribosomes were pooled and pelleted by ultracentrifugation (Ti 70 rotor, 36,600 rpm, 5 h) and dissolved in 70S buffer by gentle shaking.

### Ribosome binding assay Saccharomyces cerevisiae

For ribosome binding assays 1 pmol of ribosomes were incubated with 20 pmol His_6_-Loc1p in a total volume of 25 µl in binding buffer (20 mM HEPES pH 7.5; 150 mM KOAc; 10 mM Mg(OAC)_2_; 1 mM DTT) for 15 min at room temperature (RT) and 10 min at 4 °C. The binding reaction was loaded onto 625 µl of 0.75 M sucrose in binding buffer and ultracentrifuged in a Ti 55 rotor for 2.5 h at 39,000 rpm. The tube was flash frozen in liquid nitrogen. The sample was divided into two parts (supernatant and pellet) that were TCA precipitated and analyzed by 13% SDS PAGE stained by Sypro Orange (Sigma-Aldrich). Experiments were performed at least twice (n ≥ 2).

### Ribosome binding assay *Mycobacterium smegmatis*

For ribosome binding assays, 130 nM ribosomes were incubated with 1.3 µM His_6_ Loc1p in a total volume of 29 µl in binding buffer (20 mM HEPES pH 7.4; 200 mM NH_4_Cl; 6 mM MgCl_2_; 3 mM beta-mercaptoethanol) for 15 min at RT and 10 min at 4° C. The binding reaction was loaded onto 600 µl of 0.75 M sucrose in binding buffer and ultracentrifuged for 225 min at 33,000 rpm in a SW-55 rotor. The tubes were flash-frozen in liquid nitrogen. The sample was divided into two parts (supernatant and pellet) that were TCA precipitated and analyzed by 15% SDS PAGE stained with Sypro Orange. Experiments were performed at least twice (n ≥ 2).

### Purification of yeast tRNA

Wild-type (S288c) and *loc1Δ* cells were grown in YPD or YPAD media to the early log phase. Cells were harvested (10 min at 4,500 rpm), resuspended in buffer (10 mM Mg(OAc)_2_; 50 mM NaOAc; 150 mM NaCl; pH adjusted to 4.5), and again harvested (10 min 4,500 rpm) before they were flash frozen in liquid nitrogen. For tRNA purification cells were resuspended in buffer (10 mM Mg(OAc)_2_; 50 mM Tris; pH adjusted to 7.5) and an equal amount of phenol was added. The suspension was incubated for 1 h at 65 °C interrupted by vigorous vortexing every 10 min. After centrifugation (10,000 x g, 1 h at 20 °C) an equal volume of phenol was added to the supernatant. Subsequently, the mixture was vortexed and again centrifuged (10,000 x g, 30 min at 20 °C). The supernatant was precipitated with 0.1 volume 5 M NaCl and 2 volumes 100% ethanol at -20 °C overnight. The extracted RNA was resuspended in buffer A (40 mM Na_2_HPO_4_ pH 7) and bound to a MonoQ anion-exchange column. A gradient to 100% buffer B (40 mM Na_2_HPO_4_ pH 7; 1 M NaCl) was applied and fractions containing pure tRNAs, as assessed by denaturing PAGE, were pooled and precipitated. For mass-spectrometric analysis pellets were resuspended in water and hydrolyzed with NaOH.

### Fluorescence-based chaperone assays using molecular beacons

Fluorescence-based chaperone assays were carried out as previously described (22). In brief, assays were performed using an initial RNA concentration of 100 nM and a concentration of 75 nM of the complementary, unlabeled RNA in a total volume of 900 µl using a FluoroMax-P fluorometer (Horiba). MBP-Loc1p and MBP were added accordingly. Curve analysis was performed using Origin 8.4. FAM fluorescence was excited at 495 nm with a slit size of 2 nm. Emission was recorded at 517 nm with a slit size of 3 nm and an integration time of 0.5 s. Experiments were performed at least three times for quantification.

### *HIV TAR* annealing assay

*TAR*-annealing assay was essentially performed as previously described (23). Briefly, 0.03 pmol radioactively labeled *TAR* (+) DNA and the equal amounts of cold *TAR* (−) DNA were incubated with the indicated protein concentrations for 5 min at 37 °C in annealing buffer (20 mM HEPES pH 7.8 NaOH; 50 mM NaCl; 2 mM MgCl_2_). A sample with buffer instead of protein served as negative control, whereas an incubation at 70 °C without protein served as positive control. The reaction was stopped by the addition of 5 µl stop solution containing 20% glycerol, 20 mM EDTA pH 8, 2% SDS, 0.25% bromophenol blue and 0.4 mg/ml yeast tRNA. Annealing experiments with Loc1p peptides Loc1p (1-20) and Loc1p (1-20)_2_ were treated with 1 µl trypsin (2 mg/ml) for 10 min at RT before the reaction was stopped. Formation of ds*TAR* DNA was resolved on 8% TBE PAGE at 4 °C. Subsequently, gels were fixed, dried and auto-radiographed. Amounts of ds*TAR* DNA were determined using an LAS phosphorimager reader and the ImageJ 1.47v software and K_m_ values calculated from at least three independent experiments.

### NMR experiments

All NMR experiments were carried out on Bruker Avance III spectrometers at field strengths corresponding to 600 and 800 MHz proton Larmor frequencies, equipped with a TCI cryogenic probe head at 278 K, if not stated otherwise. Prior NMR measurements, the RNA was snap cooled to ensure the formation of monomeric hairpin conformations. Afterwards the sample was kept on ice and measured at 278 K to stabilize the RNA secondary structure. For assignment of the imino signals a 2D NOESY was recorded, using a spectral width of 25 ppm to include the downfield region of imino signals. Water suppression was achieved by using water selective pulses and pulsed field gradients. Signal intensity changes (line broadening) and chemical shift perturbations of imino signals upon titration of different Loc1p peptides and during thermal scans were observed with 1D-^1^H-NMR spectroscopy, employing the same spectral width and water suppression schemes. To estimate the melting temperature for individual imino signals, the normalized peak intensity relative to an imino signal, which is least affected by increasing temperature in all peptide-RNA complexes and free RNA (U21), was plotted versus temperature. The melting temperature was read from the temperature at midpoint intensity. For backbone and partial side chain assignment of the ^15^N,^13^C-labeled Loc1p peptide HNCACB and HCCH-TOCSY spectra were recorded at 278 K in NMR buffer (20 mM Na_2_HPO_4_/NaH_2_PO_4_, 50 mM Na Cl, pH 7.4) at a concentration of 1.6 mM, processed with NMRPipe (24) and analyzed with CARA (http://cara.nmr.ch). For RNA Loc1p-binding studies, the RNA was titrated into ^15^N,^13^C-labeled Loc1p peptide and chemical shift perturbation were monitored by acquiring a ^1^H-^15^N-HSQC spectrum for each titration point. RNA binding occurred in the fast-exchange regime and peaks could be manually traced in the NMR spectra. NMRview (25) was used to determine the binding affinity by fitting the chemical shift perturbations to:

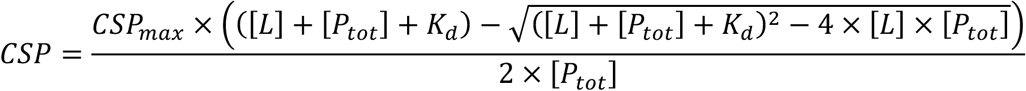

Secondary structure prediction from secondary chemical shifts (C_α_ and C_β_) was done according to Wishart and Sykes (26). 1D-^31^P-NMR has been carried out on a Bruker Avance III HD 400 MHz spectrometer equipped with a BBO ^31^P observed probehead, using a power-gated ^1^H-decoupled, ^31^P-observed pulse experiment.

### Bioinformatic analyses

#### Proteome-wide search for PUN proteins

The protein sequences of all proteins in *Saccharomyces cerevisiae* (strain ATCC 204508 / S288c), *Homo sapiens*, *Drosophila melanogaster, Schizosaccharomyces pombe* (strain 972 / ATCC 24843) and *Escherichia coli* (strain K12) were downloaded from the UniProt database (27) (UniProt IDs: UP000002311, UP000005640, UP000000803, UP000002485, UP000000625, respectively) as fasta files. We defined a PUN motif with the following regular expression: [RK](1,3)-{RK}(1,3)-[RK](2,3)-{RK}(1,3)-[RK]{1,3} where square brackets specify allowed, and curly brackets specify disallowed aa, with round brackets defining the respective number of aa. We then defined PUN proteins as proteins that (i) contain 5 or more PUN motifs and (ii) contain at least 3 PUN motifs within 150 amino acids. To account for the much shorter proteins in *E. coli*, we defined required PUN proteins in *E. coli* to contain at least 3 PUN motifs.

#### Disorderdness of PUN proteins

To assess the disorderdness of PUN proteins, we downloaded the IUPred2 scores per aa (28) using the Bioconductor R package idpr (version 1.12.0) (29) and calculated the mean IUpred2 score for each PUN protein. 500 randomly selected proteins without a single PUN motif were used for comparison. Statistical significance of the difference in the disorderdness between both groups was tested using the Wilcoxon rank-sum test.

#### Functional enrichment analysis

We performed functional enrichment analysis using Gene Ontology (GO) (30) terms for the PUN proteins with the Bioconductor R package clusterProfiler (31) together with the respective Bioconductor AnnotationDbi (version 1.64.1) (32) annotation packages for each organism (org.Sc.sgd.db version 3.18.0, org.Hs.eg.db version 3.18.0, org.Dm.eg.db version 3.18.0). For this, the UniProt IDs were transferred to Entrez gene IDs. The enrichment analysis was performed with the enrichGO() function of clusterProfiler with all proteins in the UniProt fasta file as background. We chose a P-value cut-off at 0.05 after multiple testing correction (Benjamini Hochberg). The results for all three GO subtypes – Biological Process, Cellular Component, and Molecular Function – were combined.

For the *E. coli* PUN proteins, we modified the approach because a suitable annotation package that included UniProt IDs was not available. Instead, *E. coli* Gene Ontology annotation (ecocyc) was downloaded from the GO database in .gaf format and imported with the Bioconductor R package mgsa (version 1.50.0) (33). From this, we extracted GO IDs for each protein and GO terms for each GO ID and used these as TERM2GENE and TERM2NAME parameters in the enricher() function of clusterProfiler.

#### Nuclear and nucleolar PUN proteins

To obtain all nuclear and nucleolar proteins annotated, we used the Bioconductor R package BiocSet (version 1.16.1) (34) to obtain all proteins annotated for the terms nucleus (GO:0005634) and nucleolus (GO:0005730).

#### Human orthologs of Yeast PUN proteins

We obtained orthology information from the Ensembl BioMart database (35) using the Bioconductor R package biomaRt (36). To compare whether yeast PUN proteins also exist in the human proteome only yeast proteins with one-to-one and one-to-many human orthologs were analyzed.

### Peptide Array

Loc1p tiling array was purchased from JPT peptide technologies (Berlin; Germany). N-terminally acetylated peptides with a shift of three amino acids were covalently spotted onto a beta-alanine cellulose membrane. The membrane was incubated with 0.003 nM radioactively labeled *ASH1* E3 (short) (37,38) in 10 ml buffer (20 mM Na_2_HPO_4_; 50 mM NaCl; pH 7) at 25 °C on a rotating wheel for 30 min. The membrane was washed five times with buffer for 10 min until binding was analyzed using LAS phosphorimager and Image J 1.47v software.

### Sedimentation assay

For sedimentation analysis, total RNA was purified from HeLa cells using TRIzol RNA isolation Reagent (Invitrogen). Proteins (50 µM) dissolved in an HNM buffer (20 mM HEPES, 200 mM NaCl, 2 mM MgCl_2_) were further diluted in low salt buffer of 50 mM HEPES pH 7.5 and 50 mM NaCl together with the RNA. The protein, in final concentration of 1 µM, and the RNA, in increasing concentrations (1-16 µg/ml), were incubated at 4 °C for 15 min. The samples were centrifuged at 16,100 x g at 4 °C for 1 h. The precipitate was resuspended in the same buffer with equal volume as the supernatant. Equal volumes of the pellet fraction and the supernatant were analyzed by SDS-PAGE and stained by SyproRuby. Experiments were performed at least two independent times.

### *In vitro* phase separation assays

Purified proteins were diluted in condensate buffer (20 mM Na_2_HPO_4_/NaH_2_PO_4_, pH 7.5, 150 mM NaCl, 2.5% glycerol, 1 mM DTT) to the indicated final concentrations and total HeLa cell RNA was added at the indicated RNA-to-protein mass ratios. Experimental conditions for microscopic phase separation assays underwent some evolution during the project.

Initially, samples were directly transferred to self-assembled sample chambers formed by double-sided sticky tape, taped onto an untreated glass slide and sealed with a coverslip to visualize condensates. Imaging was done by phase contrast microscopy using a 63x/1.40 Oil/Ph3 objective on an AxioCam (Zeiss, Oberkochen, Germany). For optimized experiments with round condensates, coverslips and glass slides were pre-treated with the following procedure. They were washed with purified ddH_2_O, followed by a 30 min incubation in 2% Hellmanex III. After washing 3x with purified ddH_2_O, glass slides and coverslips were incubated in 1M NaOH for 30 min, washed 3x with purified ddH_2_O, and dried in a gaseous N_2_ stream. Condensates were imaged by phase contrast microscopy using an HC PL Fluotar L 40x/0.6 PH2 objective on a Leica DMi8 (Leica, Germany). Experiments were performed at least three independent times.

### Droplet fusion analysis

Droplet fusion was recorded in mixtures containing 20 µM Loc1p and 0.5x mass ratio total HeLa RNA with the above-mentioned setup (HC PL Fluotar L 40x/0.6 PH2 objective on Leica DMi8) in movie mode. Fusion events were manually located and the aspect ratio of droplets measured in ImageJ was plotted against time. The resulting data were fitted to an exponential decay model in OriginPro 9.65. Three different fusion events were inspected.

### FRAP experiments

Fluorescence recovery after photobleaching (FRAP) was used to measure the dynamic exchange of proteins between the droplets and the surrounding protein solution using an HC PL APO 40x/1.3 OIL CS2 objective mounted on a Leica TCS SP8 inverse confocal microscope (Leica, Germany). At 40x magnification and 16x optical zoom, droplets were outlined with a circular “bleaching” overlay object. FRAP parameters were as follows: 3-4 pre-bleaching images were recorded, bleaching was done at a laser power of 100% for 1 second (HeNe laser 633 nm), and movies were recorded at a framerate of 1.95 seconds/frame. An immediate post-bleach image and several images over a total time range of 83 seconds were recorded. FRAP curves were calculated by normalizing the fluorescence signal to the background and the pre-bleach intensity. All imaging was done with the same acquisition settings (image size 512 x 512, emission wavelength 641-740 nm, scan speed 400 Hz, gain 573). FRAP experiments were repeated five times.

### Crystallization, diffraction data collection and processing

The crystallization experiments for the RNA (3’-UCUCUGGUUAGGAAACUAACUAGGGA-5’) were performed at the X-ray Crystallography Platform at Helmholtz Zentrum München. The initial crystallization screening was done at 292 K with a Mosquito (SPT Labtech) nanodrop dispenser in sitting-drop 96-well plates and commercial screens. After selecting the best hits from the screening, manual optimization was performed. The best crystals grew in 100 mM NaCl, 15 mM BaCl_2_, 40 mM Na cacodylate 7.0, 44% (v/v) MPD, and 12 mM spermine tetrahydrochloride. For the X-ray diffraction experiments, the crystals were mounted in a nylon fiber loop and flash-cooled to 100 K in liquid nitrogen. Diffraction data were collected on the SLS PXIII X06DA beamline (PSI, Villigen) and the measurements were performed at 100 K. X-ray diffraction data set for RNA was collected to 2.0 Å resolution (**Table S5**). The data set was indexed and integrated using *XDS* (39) and scaled using *SCALA* (40,41). Intensities were converted to structure-factor amplitudes using the program *TRUNCATE* (42). **Table S5** summarizes data collection and processing statistics.

### Structure determination and refinement

The structure of RNA was solved using the SAD phasing with Barium from the mother liquor, using the Auto-Rickshaw, the EMBL-Hamburg automated crystal structure determination platform (43,44). The input diffraction data were prepared and converted for use in Auto-Rickshaw using programs of the *CCP4* suite (41,45). F_A_ values were calculated using the program *SHELXC* (46). Ten heavy atom positions were located with the program *SHELXD* (46). The correct hand for the substructure was determined using the programs *ABS* (47) and *SHELXE* (46). The occupancy of all substructure atoms was refined using the program *MLPHARE* (41) and phases improved by density modification using the program *DM* (41,48). The initial model was partially built using the program *ARP/wARP* (49,50). Further model building and refinement was performed with *COOT* (51) and *REFMAC5* (52), respectively, using the maximum-likelihood target function including TLS parameters (53). The final model is characterized by R and R_free_ factors of 17.5% and 24.1% (**Table S5**). The stereochemical analysis of the final model was done in *MolProbity* (54). All crystallographic software was used from the SBGRID software bundle. Atomic coordinates and structure factors have been deposited in the Protein Data Bank under accession code 6YMC.

### Small angle X-ray scattering

Synchrotron SAXS data were collected at beamline ID14-3 (ESRF, Grenoble, France) with a wavelength of 0.931 Å. His_6_-Loc1p was measured at three different concentrations (3, 5 and 10 mg/ml) in a buffer containing 200 mM NH_4_Cl, 20 mM HEPES pH 7.5, 6 mM MgCl_2_ and 5 mM DTT with exposure times of 30 seconds each. Evaluation and processing were performed using programs from the ATSAS 2.1 software package (55). PRIMUS was used for primary data analysis and to exclude potential sample aggregation during the measurement. Based on Guinier plots, very similar radii of gyration (R_g_) between 4.6 and 4.8 nm were calculated for the three different Loc1p concentrations. Buffer-corrected profiles of low, medium and high Loc1p concentrations were used to calculate individual Kratky plots. As a reference, bovine serum albumin (5.7 mg/ml) was measured in a buffer containing 50 mM HEPES (pH 7.5) and 2 mM DTT.

### Reagents

Commercially available enzymes Thrombin (112374, Sigma-Aldrich), DNase I (A3778, AppliChem) and Trypsin (T8003, Sigma-Aldrich) were purchased. The peptides for the peptide tiling array were purchased from JPT Peptide Technologies. The PureLink RNAMini Kit (12183020, ThermoFisher) was used for the isolation of total HeLa RNA.

### Biological Resources

*Escherichia coli* strains BL21 (DE3) Star, ER2566, and BL21-Gold (DE3) pRARE were used for protein expression (see details below). HeLa cells were used for total RNA isolation. *Saccharomyces cerevisiae* wild type (MATa, his3ι11, leu2ι10, met15ι10, ura3ι10) and *loc1*ι1 (MATa, ade21, can1100, his311,15, leu23,112, trp11, ura3, GAL, psi+, loc1::natNT2) strains were used.

### Statistical Analyses

Information on the respective statistical method and corresponding sample sizes are given in the figure legends and the corresponding methods section.

### Novel Programs, Software, Algorithms

The code for the computational analyses is available at https://doi.org/10.5281/zenodo.13348876

### Web Sites/Data Base Referencing (micro-array data)

http://puma.princeton.edu/: experiment set No. 7342 https://www.ncbi.nlm.nih.gov/geo/: experiment set No. GSE273374.

## Results

### *loc1Δ* cells show growth defects and depletion of ribosomes

In order to dissect the function of Loc1p, we first compared the growth rate of a *loc1Δ* strain in suspension culture with that of a wild-type *S. cerevisiae* culture. For the *loc1Δ* strain, we observed a 2-fold increased doubling time compared to wild-type cells (**Figure 1A**). This finding is consistent with previously reported ribosome profiles showing that 80S ribosomes, half-mer ribosomes, and 60S ribosomal particles are less abundant in *loc1Δ* cells compared to wild-type (4,14). Considering the central importance of ribosomes for cellular functions, these data suggest that the reduced amount of ribosomes is responsible for the observed slow-growth phenotype of the *loc1Δ* strain.

**Figure 1:**
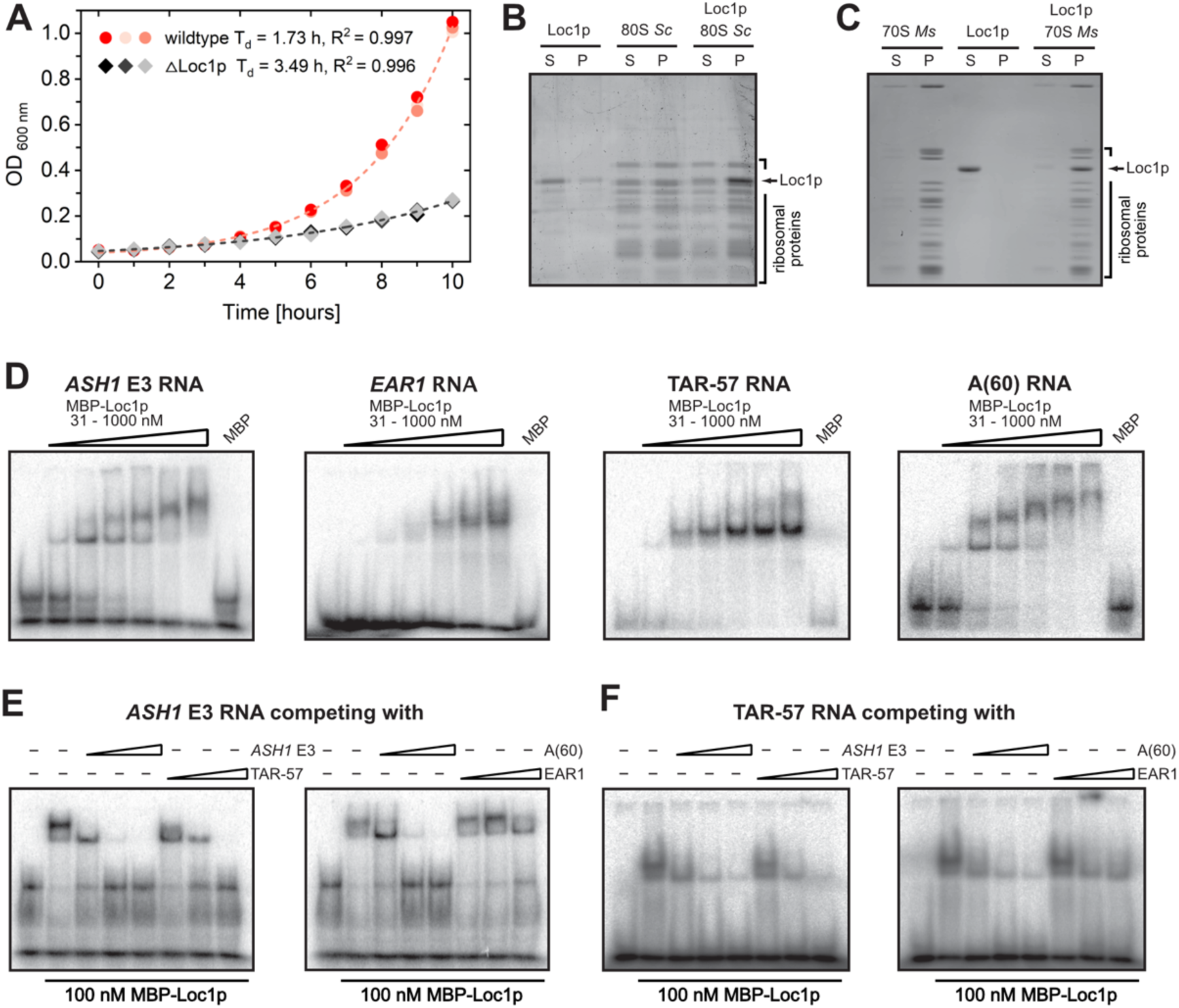
Loc1p binds directly to ribosomes and unspecifically to RNA. **(A)** Comparison of doubling time of wt (S288C) and *loc1Δ* strains. *loc1Δ* cells show a severe growth delay (n=3). **(B** and **C)** Analytical ultracentrifugation of *S. cerevisiae* (B) and *M. smegmatis* (C) ribosomes and His_6_-tagged Loc1p reveals a co-enrichment of Loc1p with ribosomes in the pellet. In absence of ribosomes, Loc1p is mainly found in the supernatant fraction of the sucrose cushion. **(D)** EMSAs with Loc1p and radioactively labeled *ASH1* E3 localization element (LE), *EAR1* LE, *HIV-TAR* stem-loop and Poly-A_60_ RNA reveal unspecific binding in the nanomolar range. **(E-F)** Competition EMSAs with 100 nM MBP-Loc1p, 5 nM radioactively labeled RNA and 10-, 100-, and 500-fold excess of cold competitor RNA. Experiments show that *ASH1* E3, *TAR*-57 and Poly-A_60_ can compete for Loc1p interaction with *ASH1* E3 **(E)** or *TAR*-57 **(F)**.

### Loc1p associates with rRNAs of the large ribosomal subunit

In order to assess the cellular binding targets of Loc1p, we immunoprecipitated TAP-tagged Loc1p expressed in yeast under the endogenous promoter and identified the co-purified RNAs by cDNA microarrays. Amongst the co-purified RNAs, the class of rRNAs was highly enriched (**Figure S1A**, **Tables S2** and **S3**), suggesting that this nucleolar RNA-binding protein preferentially binds to rRNAs. In addition, open reading frames located in proximity of rDNA fragments were amongst the most abundant Loc1p targets. Together with the previously reported defects in early rRNA processing in *loc1Δ* cells and the nucleolar localization of Loc1p (15), these data strongly suggest a direct role of Loc1p in early rRNA processing. Nevertheless, Loc1p was also associated with mRNAs (379 protein-coding genes at 1% FDR), a significant fraction of them coding for membrane proteins (*P-*value < 10^-8^; 150 out of 297 membrane protein genes).

In order to assess if Loc1p directly and specifically associates with mature ribosomes, we performed ultracentrifugation using purified *Mycobacterium smegmatis* or *S. cerevisiae* ribosomes and recombinant His_6_-Loc1p. When centrifuged separately, ribosomes sedimented in the pellet and the majority of Loc1p remained in the supernatant. However, when recombinant His_6_-Loc1p was mixed with ribosomes, Loc1p became enriched in the pellet (**Figure 1B,C**), indicating a direct interaction between Loc1p and assembled ribosomes. The interaction with both, pro-and eukaryotic ribosomes suggests unspecific binding to rRNA or ribosomal proteins and a rather general mode of ribosome association.

### Loc1p is an unspecific nucleic acid-binding protein

In order to systematically assess the specificity of Loc1p for its target RNAs, we performed electrophoretic mobility shift assays (EMSAs) with radioactively labeled RNAs. Loc1p bound with similar affinities to its previously reported RNA target sites in the *ASH1* and *EAR1* mRNAs (38,56), to the *HIV-TAR* control RNA, and even to poly-A RNA (**Figure 1D**). Binding to the E3 element of the *ASH1* RNA could be competed for by an excess of unlabeled *ASH1*, *HIV-TAR* or by poly-A RNAs, but not by the *EAR1* RNA (**Figure 1E**). Also, Loc1p binding to the *HIV-TAR* RNA was successfully competed for by unlabeled *ASH1*, *HIV-TAR*, poly-A, and *EAR1* RNAs (**Figure 1F**). In summary, we observe unspecific RNA binding by Loc1p that exhibits only minor differences in binding affinities in competition experiments.

In order to find out whether Loc1p preferentially binds to single-stranded (ss) or to double-stranded (ds) DNA or RNA, we performed EMSAs with DNA and RNA. In a first control experiment, mobility shifts by Loc1p were observed for both ssRNA and ssDNA with similar binding affinities (**Figure S1B,C**). Next, we performed EMSAs with in-gel FRET to compare Loc1p binding to dsRNA and to an RNA-DNA hybrid. In-gel FRET analysis after native PAGE confirmed binding and showed that both strands remained hybridized upon Loc1p binding (**Figure S1D,E**). In summary, we conclude that Loc1p binds indiscriminately to ssRNA, ssDNA, dsRNA, as well as DNA-RNA hybrids. Together with the observed lack of sequence preference (**Figure 1D-F**), these findings show that Loc1p interacts with general features of nucleic acids, such as their phosphate-ribose backbone.

### Loc1p acts as RNA-folding catalyst

How could the observed unspecific RNA binding be important for ribosome biogenesis? An intriguing possibility is that binding of Loc1p to RNA influences its conformation and folding required for proper RNP biogenesis in the RNA-crowded nucleolus. To assess this possibility, we performed *in vitro* RNA-chaperone experiments with a 27-base stem-loop labeled at its 5’-end with a fluorescent dye and at its 3’-end with a quencher molecule (22,57). Separation of annealed strands results in less quenching and thus higher fluorescence intensity. When increasing amounts of Loc1p were titrated to this RNA stem-loop no significant increase in fluorescent signal was observed (**Figure 2A**), even at concentrations several fold higher than required for efficient dsRNA binding (**Figure 1D**). From this, we conclude that Loc1p does not melt dsRNA.

**Figure 2:**
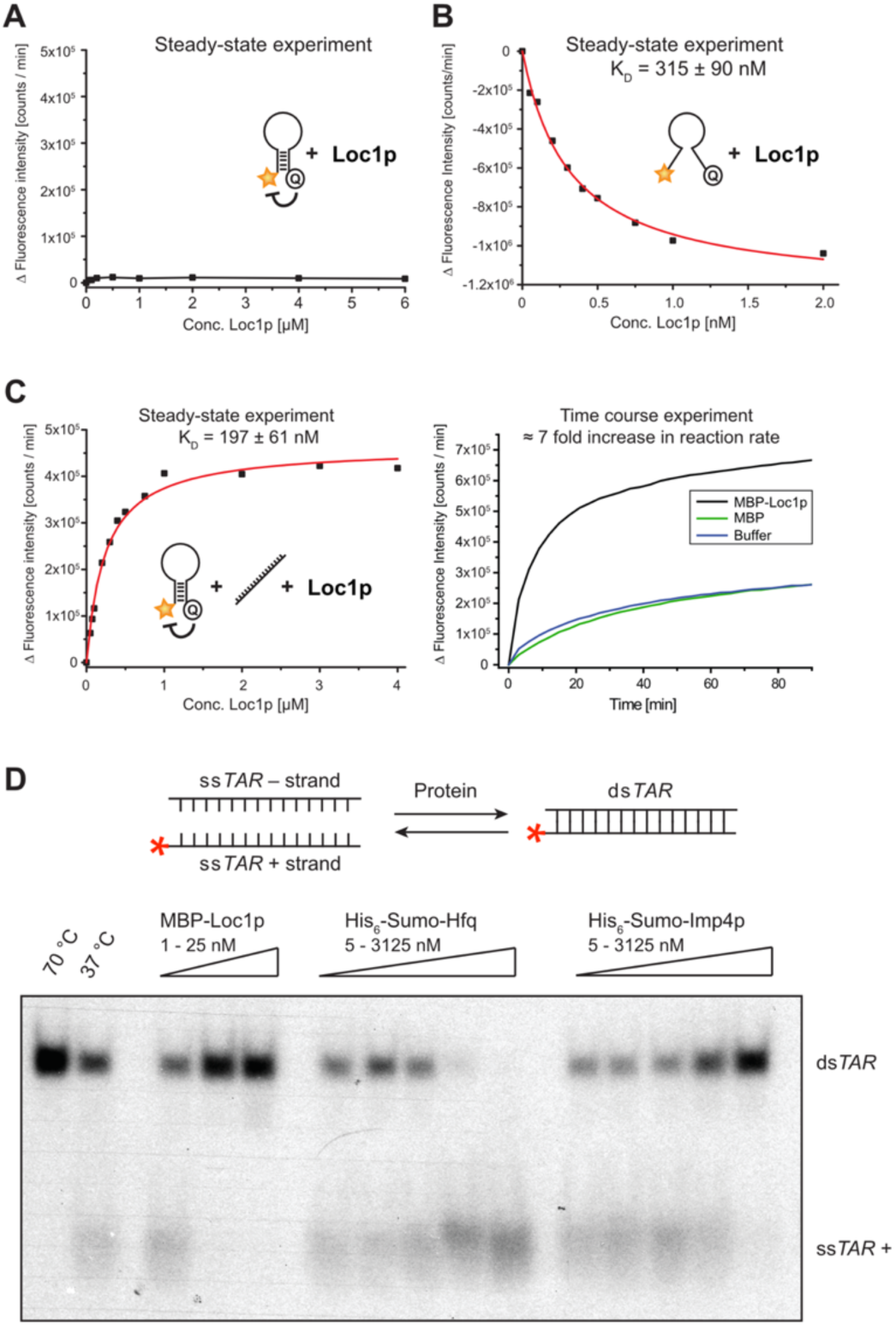
Loc1p induces conformational changes in nucleic acids. **(A)** Fluorescence intensity of a stem-loop molecular beacon (MB-D16) is not changed upon titration with MBP-Loc1p, indicating that Loc1p is unable to change the secondary structure of this RNA. **(B)** Fluorescence intensity of a single-stranded molecular beacon (MB-A5) is decreased upon titration of Loc1p, indicating the induction of conformational changes. **(C)** In presence of antisense RNA (anti-MB-D16), titration of Loc1p changes the fluorescence intensity of the stem-loop shown in (A). This suggests that Loc1p promotes the formation of the energetically favored structure. Left: For steady-state experiments the molecular beacon was pre-incubated with antisense RNA before Loc1p was added. Right: In time-course experiments Loc1p, MBP or buffer alone was pre-incubated with the molecular beacon prior to addition of antisense RNA. Addition of 1 µM Loc1p increases the reaction rate about seven-fold, suggesting that Loc1p promotes the formation of the energetically most favored state. **(D)** Comparative *TAR*-annealing assay showing that MBP-Loc1p and Imp4p promote annealing of dsDNA while Hfq displaces ds*TAR* DNA under these experimental conditions. Radioactively labeled *TAR* (+) DNA was incubated with *TAR* (−) DNA at 37 °C and with MBP-Loc1p, SUMO-Hfq, SUMO-Imp4p or buffer (negative control, 37 °C). As positive control, the sample was incubated at 70 °C for heat-induced annealing. While 5 nM of MBP-Loc1p is sufficient for complete annealing, Imp4p requires more than 3 µM to reach a comparable activity. All experiments were performed independently at least three times.

In contrast, when an ssRNA oligonucleotide that fails to form stable double-strands was used, the addition of Loc1p resulted in a concentration-dependent quenching of the fluorescence (**Figure 2B**). This indicated that Loc1p brings 5’ and 3’ ends of the ssRNA in closer proximity. Finally, we tested the ability of Loc1p to promote or inhibit the transition from an intermediate state of RNA folding to a more stable, energetically favored fold. For this, we used the above-described stem-loop RNA (**Figure 2A**) and added unlabeled, fully complementary RNA (**Figure 2C**, left panel). Since the unlabeled, complementary oligonucleotide hybridizes over its entire length with the hairpin sequence, the double-stranded heterodimer should be the energetically preferred folding state. However, to reach this state the energetic barrier of melting the intramolecular stem interactions has to be overcome. In the absence of protein or the presence of maltose binding protein (MBP) containing numerous positive surface charges, this transition occurred at a rather slow rate. In contrast, in the presence of Loc1p we observed an about seven-fold increased reaction rate towards the formation of the energetically favored dsRNA (**Figure 2C**, right panel). Thus, similar to a catalytic reaction, Loc1p accelerates the transition of folded RNAs toward their thermodynamically most favored state.

### Comparison of Loc1p with other RNA folding catalysts

In order to compare the activity of Loc1p as folding catalyst with previously reported RNA chaperones, we performed comparative annealing assays with Loc1p, Imp4p, and Hfq. Imp4p is an RNA chaperone required in the early biogenesis of eukaryotic ribosomes (58,59). Hfq is a bacterial protein capable of RNA annealing or strand displacement, depending on the specific experimental condition (60).

For comparison of catalytic activities, we used a previously described *HIV-TAR* annealing assay (23). Radioactively labeled *TAR* plus strand (*TAR* +) and unlabeled minus strand (*TAR* −) DNA were incubated with the different proteins at 37 °C and the formation of double-strands was assessed by native PAGE. In this assay, Loc1p showed by far the strongest annealing activity of all tested chaperones (**Figure 2D**). Already 5 nM of Loc1p were sufficient to completely anneal *TAR* + and *TAR* − DNA strands. Imp4p also annealed these oligonucleotides, but a protein concentration of more than 3 µM was necessary for complete annealing. A similar amount of Hfq had to be used to induce a complete strand separation in this assay. Thus, under these experimental conditions, Loc1p was the most potent nucleic acid annealing catalyst among the tested chaperones. These findings further support our observations from **Figures 2B-C**.

### Loc1p lacks recognizable domain folds and remains unfolded upon RNA binding

Although Loc1p is a potent RNA binder, it lacks any recognizable similarity to known RNA-binding domains. The Kratky plot of small angle X-ray scattering (SAXS) data of Loc1p at different concentrations showed that it is intrinsically disordered and lacks a globular fold (**Figure 3A**). For a more detailed assessment, we performed solution NMR experiments with ^1^H-^15^N correlation spectra of ^15^N-labeled Loc1p (**Figure 3B**). The spectrum showed poor signal dispersion, confirming that Loc1p is largely unstructured in absence of a binding partner.

**Figure 3:**
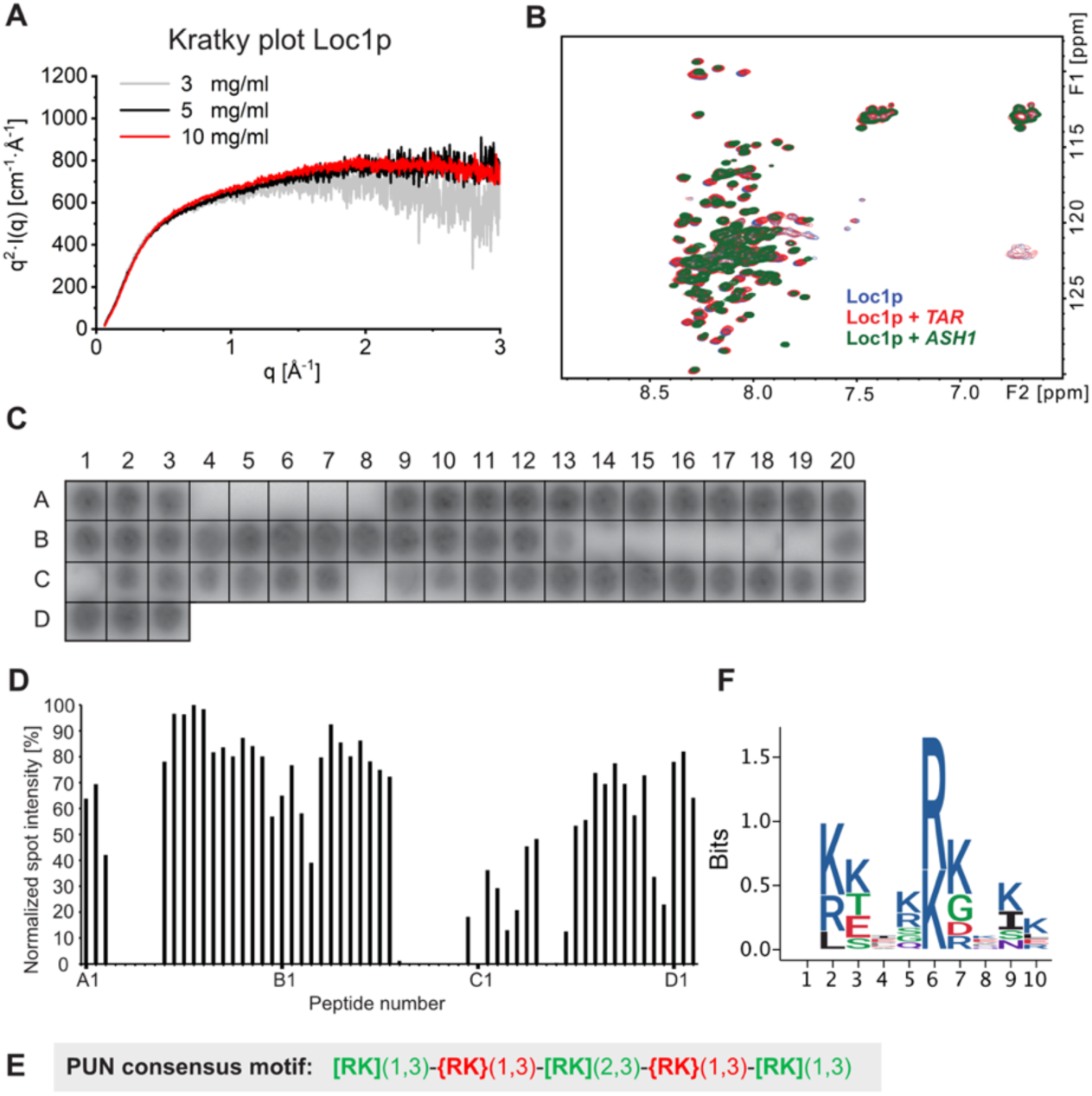
RNA binding of the intrinsically unstructured Loc1p is mediated by distinct motifs within the Loc1p sequence. **(A)** Kratky plots of SAXS measurements at three Loc1p concentrations indicate that the protein is largely unfolded. The scattering intensity (as I(q)xq^2^) is plotted against the scattering vector q. Unfolded/highly flexible proteins are typically characterized by a plateau at higher q values. **(B)** ^1^H-^15^N HSQC spectra of Loc1p alone (blue) and in the presence of *TAR* RNA (red) and the *ASH1* E1 RNA (green). Minor signal shifts indicate that the interaction with RNA does not induce a secondary structure fold in large parts of Loc1p. **(C)** Binding experiments of radioactively labeled *ASH1* E3 (short) RNA to a Loc1p peptide array revealed binding of a large portion of the spotted peptides to the RNA. Peptides had a length of 20 amino acids and an offset of 3 amino acids. **(D)** Bar-graph depicting RNA binding intensities of each peptide. Signals from the peptide array shown in (C) were quantified with Image J and normalized against the best binding peptide (A12 set to 100%; n=1). Large sequence stretches of Loc1p show binding to *ASH1* E3 (short) RNA. **(E)** The consensus sequence of the PUN motif is characterized by three repetitive units of 1-3 positively charged residues ([RK](1,3) in green) interspaced by 1-3 residues that are not positively charged ({RK}(1,3) in red). **(F)** One-letter-code sequence logo integrating all eight PUN motifs of Loc1p (amino acid chemistry: blue, basic; red, acidic; black, hydrophobic; purple, neutral; green, polar). The amino-acid position is plotted against the information content of each position in bits.

In order to assess whether RNA binding induces folding of Loc1p, we titrated different RNAs to Loc1p and monitored NMR spectral changes. Addition of either a 16 nt *HIV-TAR* stem-loop RNA or a confirmed target of Loc1p, the 49 nt E3 stem-loop of the *ASH1* RNA, to ^15^N-labeled Loc1p yielded chemical shift perturbations to some extent, but no significantly increased spectral dispersion (**Figure 3B**). Thus, Loc1p remains largely unstructured even when bound to the tested RNAs.

### A repetitive motif found throughout Loc1p mediates RNA binding

To characterize the RNA interaction within the unstructured Loc1p, we performed RNA binding experiments using a tiling array with 20 residue-long peptides of the 204 amino acid (aa)-long Loc1p, each shifted by 3 aa. In these experiments, 50 out of 63 peptides along the entire sequence of Loc1p showed significant binding to a radioactively labeled *ASH1* E3 RNA (**Figure 3C,D**, **Figure S2**) indicating the presence of several RNA-binding motifs along the protein sequence.

Analyses of the correlation between peptide sequences of Loc1p and RNA-binding activity (**Figure 3C,D**, **Figures S2** and **S3A**) identified a defined pattern of positively charged amino acids with short spacers that bound RNA (**Figure 3E**). The spacers were characterized by the absence of positively charged residues and the presence of one or more acidic residues. This pattern of alternating motifs and spacers occurred eight times in Loc1p (**Figure S3**). Motif-containing peptides that had no or only a single negatively charged side chain, such as peptides A9, A10, A11, A14, A15, B5, B6, B7, B8, or C15, tended to show the highest binding signal. In contrast, peptides with at least three negatively charged residues, like A4, A5, A7, A8, or B14-C12, showed on average weak or no binding (**Figure 3C,D**, **Figure S2**). Thus, the RNA-binding activity of these repetitive motifs in Loc1p requires positively charged amino acids and tolerates only few negatively charged residues.

### A single RNA-binding motif is sufficient to stabilize double-stranded RNA

In order to understand the molecular details of the function of this RNA-binding motif, we performed structural analyses of the first motif in Loc1p consisting of aa 3-15 (Loc1p (3-15)) and a double-stranded RNA sequence with eleven base pairs fused to a minimal tetraloop (dsRNA tetraloop, **Figure S4A**). The crystal structure of this dsRNA tetraloop alone at 2.0 Å resolution revealed a complete base-pairing in the stem region as well as a highly coordinated loop (**Figure 4A** and **Table S5**).

**Figure 4:**
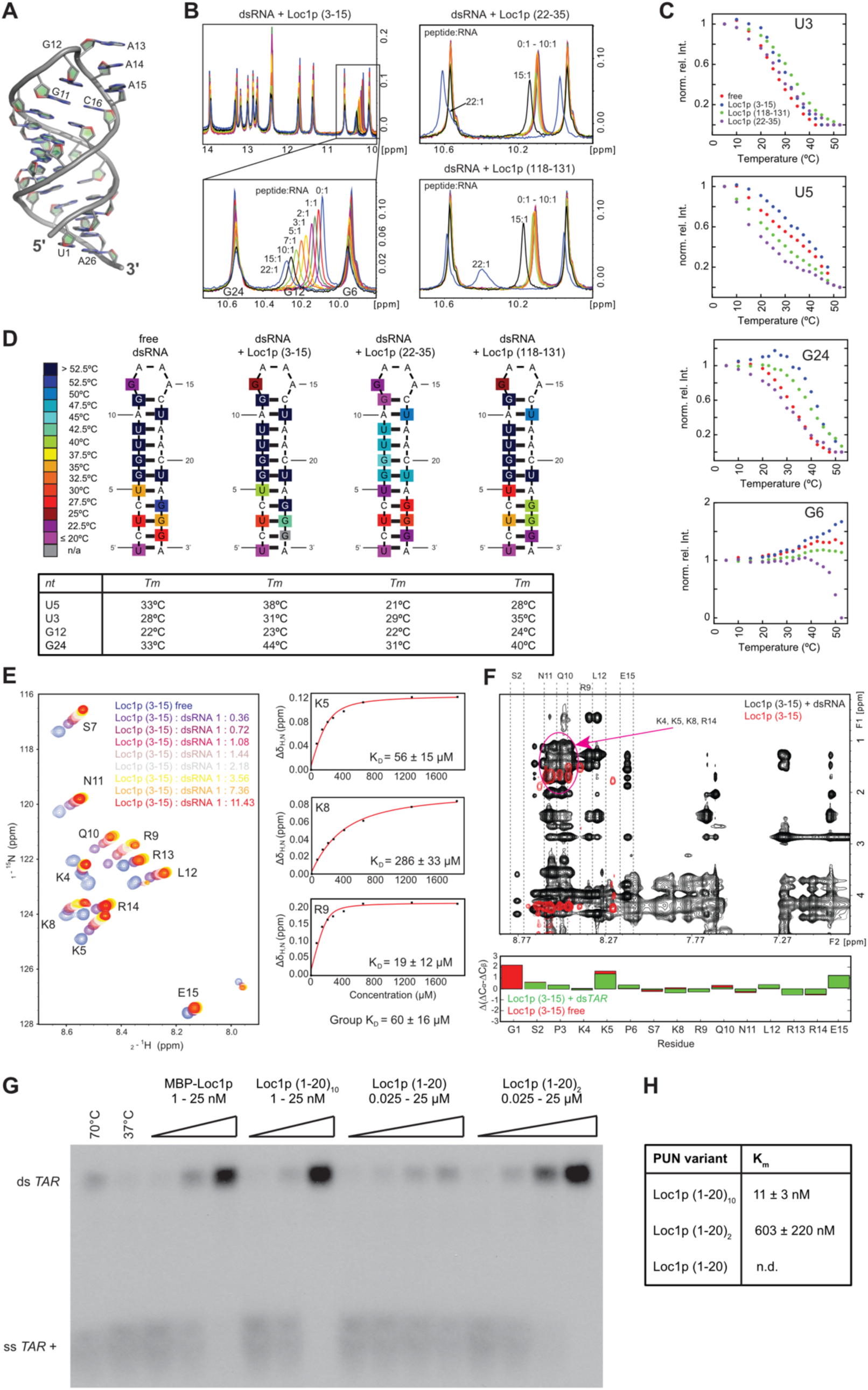
Structure analysis of RNA binding and assessment of annealing activity of PUN motifs. **(A)** X-ray structure of the dsRNA tetraloop highlighting the coordinated GAAA minimal tetraloop (G12 - A15) and the base-paired stem. **(B)** Titration of the dsRNA with different Loc1p derived peptides. Whereas the titration of Loc1p (3-15) shifts the signal of G12 in a continuous manner, the two peptides lacking the RNA binding motif Loc1p (118-131) and Loc1p (22-35) do not bind until very high, 15-fold excess, where peaks start to display sudden large shifts. **(C)** Temperature dependence of imino NMR signals of unbound dsRNA and dsRNA bound to different Loc1p derived peptides at the highest molar excess (22:1) shown in (B). The decrease of imino-signal intensities (shown: U3, U5, G6, G24) was used to determine the stabilizing effect of the three different peptides titrated in (B) in temperature scans. Whereas Loc1p (3-15) (blue dots) has a stabilizing effect on the hairpin RNA (free RNA: red dots), the control peptides Loc1p (118-131) (green dots) and Loc1p (22-35) (purple dots) have an overall destabilizing effect. The only exception is a slight stabilizing effect for U3 by Loc1p (118-131). **(D)** Heat map, illustrating the temperature-dependent stabilization of the end of the stem upon peptide binding. In contrast, Loc1p (118-131) and especially Loc1p (22-35) destabilize the ends of the stem and the loop regions. Loc1p (22-35) has even a destabilizing effect on the central stem region. The estimated melting temperatures for the four most affected base pairs are given in the table below. **(E)** ^1^H,^15^N-HSQC spectrum showing the titration of dsRNA to ^15^N, ^13^C-labeled Loc1p peptide. Saturation is reached at a 3 to 4-fold excess of dsRNA tetraloop. To the right three examples are shown of fitted binding curves from chemical shift perturbations. **(F)** 2D NOESY spectra overlay of unbound Loc1p peptide (red) and dsRNA-bound Loc1p (black). In the free state, the Loc1p peptide is flexible, as only intraresidual and sequential H_N_-H_α-1_ NOE cross peaks are observed. Upon binding to dsRNA, additional NOEs can be observed, especially non-sequential and interresidual NOEs. The dashed lines indicate strips from the H_N_ region of indicated residues to other protons. The purple circle indicates the strong overlap of positively charged residues (K4, K5, K8, R14), which prevents full assignment of peptide-side chains. Secondary chemical shifts (below the spectrum) clearly show that no secondary structure is formed upon RNA binding. **(G)** *TAR*-annealing assay with radioactively labeled *TAR* (+) and unlabeled minus strand demonstrates that concatenation of the most N-terminal Loc1p motif is sufficient to mimic the annealing activity of wild-type Loc1p *in vitro*. Both strands were incubated with increasing concentrations of wild-type Loc1p, Loc1p (1-20)_10_, Loc1p (1-20)_2_, Loc1p (1-20) or buffer (negative control: NC) at 37°C. Heat-induced displacement at 70 °C and in absence of Loc1p served as positive control (PC). Formation of ds*TAR* was analyzed by 8% native TBE PAGE. Wild-type Loc1p has an annealing activity comparable to an artificial protein bearing ten Loc1p motifs (Loc1p (1-20)**_10_**). **(H)** Comparison of annealing activities of sequences bearing one, two, and ten Loc1p motifs. For Loc1p (1-20) annealing activity was not determined because even with 50 µM no saturation was reached. According to the intensities of annealed ds*TAR* Loc1p (1-20)_2_ showed an estimated 50-100-fold higher activity than Loc1p (1-20). Furthermore Loc1p (1-20)_10_ showed an over 50-fold higher annealing activity than Loc1p (1-20)_2_. Since wild-type Loc1p showed even better reactivity that exceeded the limits of this assay, no K_m_ could be derived. Shown are quantifications from three independent experiments.

We applied NMR to characterize the binding between the first motif (Loc1p (3-15)) and the dsRNA tetraloop. The assignment of imino cross-correlation signals of the RNA alone confirmed the hairpin conformation of the dsRNA tetraloop in solution (**Figure S4A**). Next, we titrated Loc1p (3-15) with up to a 22-fold molar excess to the dsRNA tetraloop and recorded 1D spectra of nucleotide-imino signals. Increasing concentrations of Loc1p (3-15) induced a continuous signal shift of the G12 imino signal (**Figure 4B**, left). This indicated a strong interaction of the peptide in this region of the RNA, whereas other signals were only moderately affected. Chemical shifts of the imino signals of the RNA remained largely unchanged upon titration of the peptide, showing that the overall secondary structure of a dsRNA is not altered upon binding. This observation is consistent with the effect of full-length Loc1p in chaperone assays (**Figure 2A**).

When titrating the peptides Loc1p (22-35) and Loc1p (118-131), which lacked the RNA-binding motifs (**Figure S3A,C**), no NMR-chemical shift changes were observed even at 10-fold excess of peptides over RNA. Only very high excess of 15-and 22-fold resulted in larger shifts of the G12 imino signal (**Figure 4B**, right). Considering the high molar excess with peptide concentrations of up to 2.2 mM, this interaction is significantly weaker and may reflect a more unspecific interaction with these peptides. This interpretation is consistent with results from the tiling array where both peptides failed to bind RNA (peptides A7 and B19 in **Figure 3C,D** and **Figure S2**).

### NMR studies confirm that peptide binding stabilizes RNA-secondary structures

In order to understand the specific contribution of the peptide motif, we recorded 1D ^1^H spectra of RNA-imino signals at increasing temperatures in absence and presence of Loc1p peptides. The normalized signal intensities of respective imino signals were used to generate melting curves (**Figure 4C**, **Figure S4C,D**). In presence of the first motif Loc1p (3-15) the melting curves of the imino signals of U3, U5, and G24 were shifted to higher temperatures, indicating a stabilization of the stem-loop. In summary, melting points of imino signals at the 5’and 3’region, and in the loop region were increased (**Figure 4D**). In contrast, the control peptides Loc1p (22–35) and Loc1p (118-131) led to a decrease in melting points of the majority of imino signals (**Figure 4D**), indicating destabilization of the dsRNA. In conclusion, these data indicate that the positively charged RNA binding motif identified contributes to the stabilization of RNA-secondary structures. Of note, titration of high concentrations of each of the three peptides resulted in resonance shifts in 1D-^31^P-NMR experiments. However, Loc1p (3-15) induced the strongest chemical shift perturbations (**Figure S4B**), consistent with its interaction with phosphates in the phosphoribose backbone.

To analyze how the RNA interaction affects the conformation of an RNA-binding peptide, we performed NMR experiments with ^15^N and ^13^C isotope-labeled Loc1p (3-15). After side-chain assignment, we performed a titration with the dsRNA tetraloop and recorded ^1^H-^15^N heteronuclear single quantum coherence (HSQC) spectra (**Figure 4E**, left). Upon addition of RNA, we observed shifts of all peptide signals. Analyzing the chemical shift changes as a function of RNA concentration yielded K_D_ values in the range from 19 µM to 286 µM (**Figure 4E**, right). Nuclear Overhauser effect spectroscopy (NOESY) of the motif in absence of RNA showed only few intraresidual and sequential H_N_-Hα_1_ cross-signals, indicating that the peptide is flexible and not structured (**Figure 4F**, red, **Figure S4E**). In the presence of RNA, the NOESY spectrum contained significantly more signals (**Figure 4F**, black), with a particular increase of intraresidual signals. For instance, the methyl group of leucine 12 shows an NOE to the amide proton of arginine 9, indicating their close proximity. The increase in signal quantity indicates that amino acid side chains of the dsRNA-bound motif have less rotational freedom and hence are more rigid than the free peptide. Analysis of ^13^C secondary chemical shifts (**Figure 4F**, bottom), however, clearly showed that the motif does not adopt a defined secondary structure when bound to the RNA. In summary, these results indicate that Loc1p (3–15) binds to the dsRNA tetraloop with a more rigid conformation without a specific secondary structure.

Because of their functional and structural properties, we named the RNA-binding elements of Loc1p positively charged, unstructured nucleic acid binding (PUN) motifs.

### Multiple <u>p</u>ositively charged, <u>u</u>nstructured <u>n</u>ucleic acid binding (PUN) motifs cooperate for RNA annealing

Since in the tiling array, each PUN motif-containing peptide showed RNA binding, a single PUN motif might already be sufficient for RNA-annealing activity. In order to test this possibility, we compared the chaperone activity of wild-type Loc1p (204 aa with eight motifs) with a peptide bearing only the most N-terminal PUN motif (Loc1p (1-20); **Figure S3A,C**). We also used peptides containing two (Loc1p (1-20)_2_) or ten (Loc1p (1-20)_10_) of this N-terminal PUN motif. Whereas a single motif (Loc1p (1-20)) only showed weak annealing activity at the highest concentration of 25 µM (**Figure 4G**), Loc1p (1-20)_2_ already yielded strongly improved activity (**Figure 4G,H**). Although no Michaelis-Menten constant (K_m_) could be derived for Loc1p (1-20), a comparison of its band intensities with the ones of Loc1p (1-20)_2_ indicated an about 50-fold higher activity for the latter. Loc1p (1-20)_10_ showed again a more than 50-fold improvement in activity over Loc1p (1-20)_2_ (**Figure 4G,H**). These findings suggest that the RNA-binding motifs in Loc1p act cooperatively in nucleic acid annealing. Interestingly, wild-type Loc1p and Loc1p (1-20)_10_ showed comparable activities in this assay (**Figure 4G**), suggesting that the concatenation of PUN motifs is sufficient to achieve wild-type annealing activity.

### Multiple PUN motifs undergo phase separation in presence of RNA

A number of recent studies demonstrated that phase separation driven by proteins or protein-nucleic acid complexes is a prevalent organization principle of macromolecular assemblies in cells (61). Phase separation is thought to be the driving force for establishing functionally distinct membrane-less organelles such as stress granules, processing (P) bodies, and even the nucleolus (62–64). Since Loc1p is located in the nucleolus and mainly consists of low-complexity regions that could potentially promote phase separation, we tested whether Loc1p with its PUN motifs is able to undergo RNA-mediated phase separation. We first mixed increasing amounts of recombinant Loc1p with 0.5x mass ratio of total HeLa cell RNA and observed formation of microscopically visible condensates at Loc1p concentrations of 5 µM or higher (**Figure 5A**). The number and size of the condensates were clearly dependent on the Loc1p concentration, while 30 µM Loc1p without RNA did not result in phase separation. For further studies, we used an improved protocol to obtain condensates with perfectly round shapes with typical diameters between 1 and 5 µm (**Figure 5B**) at a concentration of 20 µM Loc1p and 0.5x mass ratio of RNA. In order to show that the observed structures were indeed droplet-like condensates and not protein-RNA precipitates, we analyzed individual droplet fusion events. Although relatively slow, condensates clearly fused and relaxed into spherical shapes after fusion (**Figure 5C** and **Movie S1**). To directly demonstrate the presence of Loc1p in these condensates, we introduced a single cysteine (Loc1p_S7C_) and labeled the protein using Cy5-maleimide (Cy5-Loc1p_S7C_). RNA/Cy5-Loc1p_S7C_ droplets showed strong fluorescence under the same conditions as wild-type Loc1p, indicating that Cy5-Loc1p_S7C_ indeed partitions into the droplets (**Figure 5D**). In addition, fluorescence recovery after photobleaching (FRAP) experiments were performed to assess the dynamics of Loc1p in these droplets. Fluorescence of Cy5-Loc1p_S7C_/RNA condensates recovered after photobleaching on average to 20-30% after 30 seconds (**Figure 5E** and **Movie S2**), demonstrating a dynamic exchange of Loc1p between the droplets and the surrounding protein solution. Both, the droplet fusion and the fluorescence recovery to around 20-30% are in line with the assumption that the observed condensates have liquid-like to gel-like properties.

**Figure 5:**
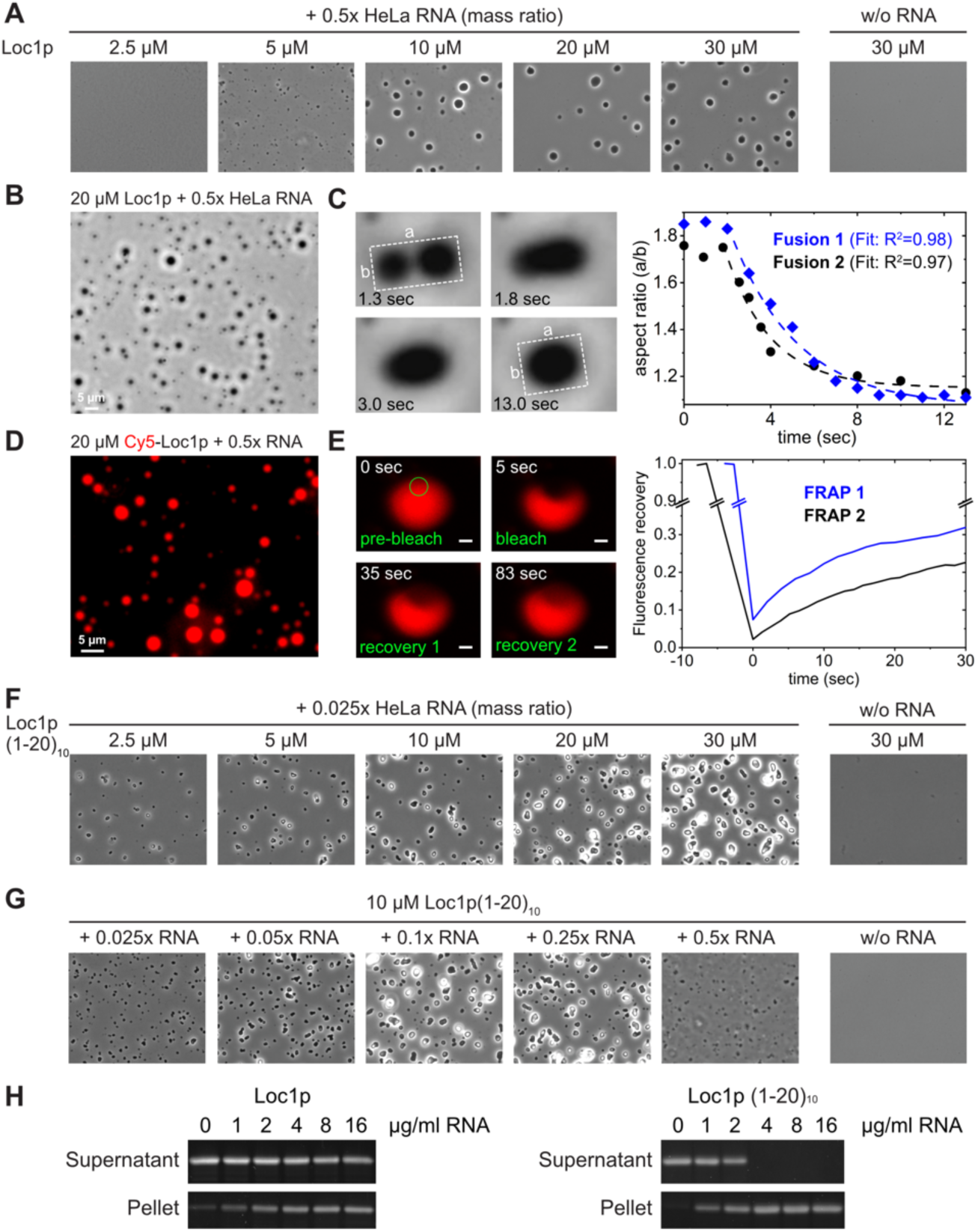
Loc1p or multiple copies of its N-terminal PUN motif form conden-sates with RNA. **(A-G)** The formation of condensates was assessed by phase contrast-and fluores-cence microscopy. **(A)** At a protein:RNA mass ratio of 0.5, phase separation was observed already at 5 µM Loc1p. **(B)** At the same protein:RNA ratio and 20 µM Loc1p, condensates had a size of 1-5 µm and showed round shapes. **(C)** The observed condensates fuse and round up after fusion, con-firming their liquid-to-gel-like properties. **(D)** Like unlabeled Loc1p, Cy5-labeled Loc1p_S7C_ promotes RNA-dependent droplet formation and partitions into the droplets. **(E)** Fluorescence of Cy5-Loc1p_S7C_/RNA droplets recover after photobleaching to 20-30% in FRAP experiments. **(F, G)** Loc1p (1-20)_10_ formed condensates already at 2.5 µM and a mass ratio of 0.025 RNA:protein and larger condensates with increasing RNA concentration. No phase separation was observed in the absence of RNA. **(H)** RNA-dependent phase separation of Loc1p and Loc1p (1-20)_10_ was confirmed by sedimentation experiments followed by SDS-PAGE stained with SyproRuby. Experiments were repeated at least once (n≥2).

As we had observed that a peptide sequence with ten consecutive PUN motifs showed high RNA chaperone activity (**Figure 4G**), we next tested whether Loc1p (1-20)_10_ was also able to trigger RNA-mediated phase separation. Indeed, Loc1p (1-20)_10_ formed visible condensates already at a concentration of 2.5 µM protein and a mass ratio of 0.025x RNA (**Figure 5F**). We further confirmed the robustness of the apparent PUN motif-mediated phase separation by providing a constant concentration of Loc1p (1-20)_10_ and increasing ratios of RNA (**Figure 5G**). In all cases, RNA-dependent phase separation was observed. In order to test these findings with an independent assay, RNA-driven phase separation of Loc1p and Loc1p (1-20)_10_ was further confirmed by centrifugation sedimentation experiments (**Figure 5H**). Since cooperative effects of multiple PUN motifs were observed in RNA chaperone assays (**Figure 4G,H**), we compared phase separation of Loc1p (1-20)_3_ or Loc1p (1-20)_5_ and RNA. Indeed, a clear correlation between the number of PUN motifs per peptide chain and their ability to mediate phase separation was observed (**Figure S5**).

To assess whether Loc1p also mediates RNA annealing and RNA-dependent phase separation on a physiological target, we performed corresponding experiments with the E3 localization element of *ASH1* mRNA. This transcript is bound co-transcriptionally by Loc1p and shuttles together through the nucleolus (7, 8, 15). Notably, formation of LLPS with an *in vitro* transcribed 28 nt long E3 (21) is observed for Loc1p concentrations higher than 5 µM but not in absence of RNA (**Figure S6A**). Furthermore, Loc1p stabilizes the energetically favored conformation of E3 in a time-course experiment with molecular beacons and antisense RNA strands (**Figure S6B**). In summary, the PUN protein Loc1p drives RNA-dependent phase separation and RNA annealing also on a physiological target transcript.

### PUN motifs are prevalent in nuclear proteins

To address whether PUN motifs are present in other proteins in yeast, we performed a computational search in the *S. cerevisiae* proteome (6,060 proteins). Based on the observation that multiple PUN motifs are required for RNA chaperone activity and phase separation, we only considered proteins with at least five PUN motifs in our search and three or more PUN motifs in close proximity (termed PUN proteins; see Methods). Intriguingly, we identified a total of 95 PUN proteins that matched both criteria, harboring up to 16 PUN motifs (**Figure 6A,B** and **Table S6**). Relative to protein length, Loc1p had the highest PUN motif density (8 PUN motifs in 204 aa) followed by Tma23p (6 PUN motifs in 211 aa), Pxr1p (6 PUN motifs in 271 aa), Efg1p, and Rpa34p (both 5 PUN motifs in 233 aa; **Figure 6B,E**). Of note, the PUN proteins have a higher degree of disorder than proteins without PUN motifs, underlining the preference of PUN motifs for disordered regions (**Figure 6C**). 84% (80 out of 95) of the PUN proteins had been reported to be located in the nucleus (**Figure 6D**) (65), and nearly half of those were also enriched within the nucleolus (37% of all PUN proteins), supporting a role of PUN proteins in ribosome biogenesis. Indeed, a functional enrichment analysis clearly showed an overrepresentation in nucleic acid and RNA-metabolic processes (**Figure 6F**), with 24 proteins being directly involved in ribosome biogenesis. Consistently, related terms such as RNP complex biogenesis, rRNA metabolic processes, and rRNA processing were also highly populated.

**Figure 6:**
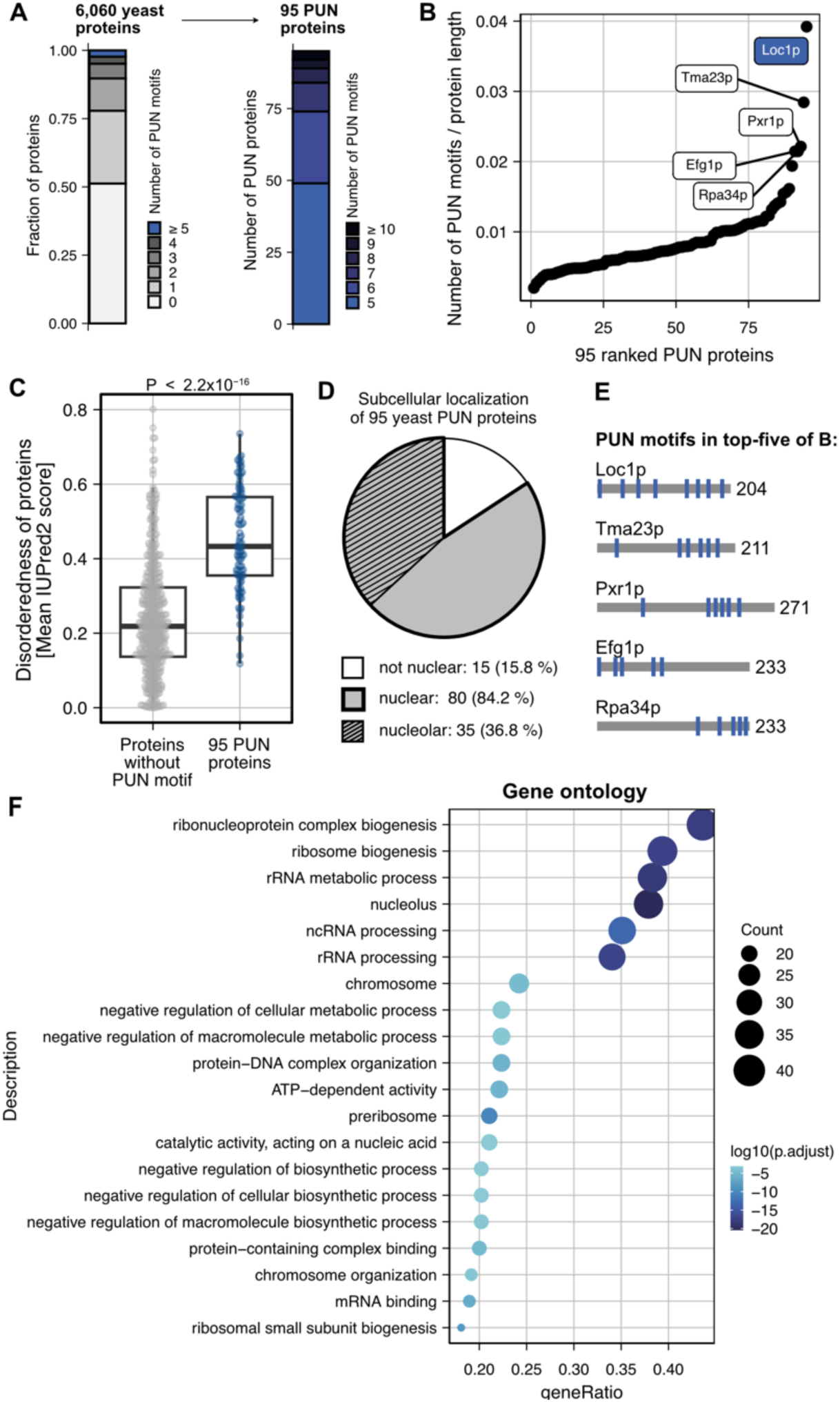
Yeast proteins containing the <u>p</u>ositively charged <u>u</u>nfolded <u>n</u>ucleic acids binding (PUN) motif identified in Loc1p are more likely to contain disordered regions, accumulate in the nucleus, and are involved in RNA metabolic processes. **(A)** Computational identification of 95 proteins from *S. cerevisiae* that contain at least five PUN motifs, including at least three within 150 amino acids. **(B)** PUN motif density in the 95 yeast PUN proteins. Loc1p has the highest density of PUN motifs followed by Tma23p, Pxr1p, Efg1p, and Rpa34p. **(C)** Yeast PUN proteins are characterized by a higher degree of disorderedness compared to proteins without PUN motifs, reflecting the preference of PUN motifs for disordered regions. P value is obtained from the Wilcoxon rank-sum test. **(D)** Analysis of the intracellular distribution of the 95 hits shows a clear preference for the nuclear (85%) and nucleolar (35%) compartments of the cell. **(E)** Distribution of PUN motifs in top-five proteins from (B). **(F)** Functional enrichment analysis (Gene Ontology, GO) of annotated processes ordered by P values. The RNA processing pathways are enriched in proteins bearing PUN motifs.

Interestingly, when we expanded our search to *Schizosaccharomyces pombe* and *Drosophila melanogaster*, we found 61 and 349 PUN proteins, respectively, which also predominantly located in the nucleus and enriched for similar functional terms (**Figure S7A,B**, **Tables S7** and **S8**). Likewise, even in the prokaryote *Escherichia coli*, proteins containing at least three PUN motifs showed significant enrichment in similar processes (**Figure S7C** and **Table S9**), suggesting functionally conserved themes associated with PUN motifs. Finally, an analysis of the human proteome yielded 500 PUN proteins with five or more PUN motifs (**Figure S8A,B,E** and **Table S10**). As for yeast, the human PUN proteins had a higher degree of disorder and were most often located in the nucleus (**Figure S8C,D**). Additionally, the human PUN proteins showed strong enrichment in nucleic acid-related processes, including mRNA processing, RNP complex biogenesis, RNA splicing, and nuclear speckles (**Figure S8F**).

To evaluate how conserved the presence of PUN motifs in proteins between distantly-related species is, we identified orthologs of all 95 yeast PUN proteins in the human proteome (**Table S11**). For approximately 36% of these proteins, no ortholog could be found while roughly 35% have one (one-to-one) or more (one-to-many) orthologs (**Figure S9A**). From these orthologous human proteins, about half fall into the definition of PUN proteins (**Figure S9B**). Further analysis showed that the location, distribution, and clustering of PUN motifs are similar between yeast PUN proteins and their human orthologs (**Figure S9C**).

### The human PUN protein SRRM1 mediates RNA-dependent phase separation

In order to experimentally assess if human PUN proteins bear the functional properties observed in Loc1p, we recombinantly expressed and purified a fragment of the human Serine/arginine repetitive matrix protein 1 (SRRM1). SRRM1 is involved in mRNA processing and splicing (66,67), rendering it an ideal candidate for comparing its potential phase separation properties to Loc1p. The chosen SRRM1 fragment (aa 151-377) had a similar size and PUN-motif content as Loc1p and was predicted to be intrinsically disordered (**Table S10**). Its N-terminal folded PWI domain was excluded from the tested fragment. To test whether this SRRM1 fragment undergoes RNA-dependent phase separation, we performed titration experiments with total HeLa cell RNA followed by microscopic analyses. At protein/RNA ratios of 0.25 and 0.5, formation of condensates was observed at protein concentrations of 2.5 µM to 30 µM SRRM1 (**Figure 7A,B**). Also, in the inverse experiment, a titration of increasing amounts of RNA at a given SRRM1 concentration of 10 µM resulted in robust condensate formation (**Figure 7C**). These findings were cross-validated in centrifugation sedimentation experiments (**Figure 7D**). In none of the experiments, condensate formation or sedimentation was observed in absence of RNA, demonstrating that SRRM1 is capable of RNA-dependent phase separation as observed for Loc1p.

**Figure 7:**
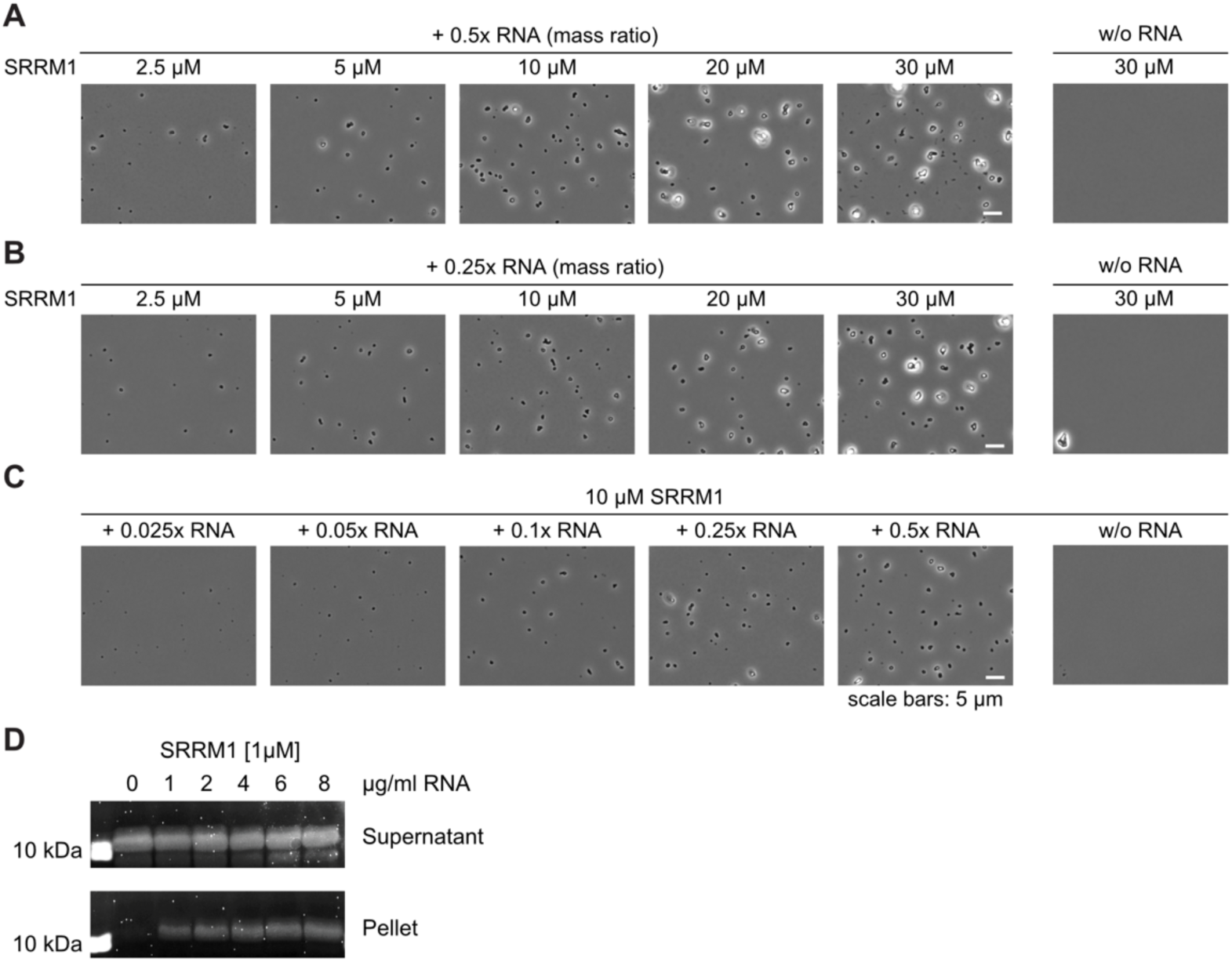
Phase separation of human SRRM1 bearing 14 PUN motifs (Table S10). SRRM1 undergoes RNA-dependent phase separation at protein concentrations of 2.5 µM or higher and at RNA:protein mass ratios of 0.5 **(A)** or 0.25 **(B)**. Titration of increasing amounts of RNA and a constant concentration of 10 µM SRRM1 showed phase separation already at the lowest RNA:protein ratio. **(D)** RNA-dependent condensation of SRRM1 was confirmed by sedimentation experiments followed by SDS-PAGE stained by SyproRuby. Experiments shown in (A-D) were repeated at least once (n≥2).

## Discussion

Loc1p was reported to be a ribosome assembly factor that assists in the synthesis of the large ribosomal subunit leading to reduced levels of 80S ribosomes and accumulation of the 35S pre-rRNA in genomic *LOC1* deletion strains (4,14,15). The presence of the aberrant 23S rRNA indicates that cleavage at the site A_3_ is unaffected, whereas co-transcriptional processing at sites A_0_, A_1_, and A_2_ is delayed. Thus, Loc1p is likely involved in early rRNA processing events. This interpretation is consistent with recent reports showing that Loc1p directly interacts with the ribosomal protein Rpl43 (13,14), as well as with other ribosome biogenesis factors and rRNA in the state E2 60S pre-ribosomal particle (12). Overall, the important role of Loc1p in ribosome biogenesis and the central importance of ribosomes for cell function are in line with the observed slow-growth phenotype of *loc1Δ* strains (**Figure 1A**) (56).

Using microarrays, we found that TAP-tagged Loc1p indeed co-purifies preferentially with rRNAs (**Figure S1A**). Furthermore, recombinant Loc1p co-sediments with mature ribosomes (**Figure 1B,C**), which suggests that Loc1p directly interacts with immature and mature ribosomes in the nucleolus. The association with ribosomes of different kingdoms of life and its unspecific binding to single-stranded and double-stranded DNA and RNA (**Figure 1D-F**, **Figure S1B-E**) suggest the recognition of general features of nucleic acids. Indeed, our ^31^P NMR experiments are consistent with a direct interaction of Loc1p with the phosphoribose backbone (**Figure S4B**). This might also explain why Loc1p is frequently found in immunoprecipitations of various RNPs (68,69). Within the cell, the nucleolar localization of Loc1p (15,65) might therefore be the main determinant of specificity for the process of ribosome biogenesis.

Using different chaperone assays, we showed that nucleic acid binding of Loc1p induces a strong annealing activity that even exceeds the activity of the well-studied U3 snoRNA–rRNA annealer Imp4p (**Figure 2**) (58,59). Subsequent NMR and SAXS analyses demonstrated that Loc1p is intrinsically disordered and lacks a globular domain fold (**Figure 3A,B**). Large parts of the unstructured Loc1p protein are involved in RNA binding (**Figure 3C,D**) through its PUN motifs (**Figure 3E**, **Figure S2** and **S3A**). While a single motif is already capable of strand annealing, the combination of two or more PUN motifs leads to a drastic increase in annealing activity (**Figure 4G,H**), indicating that the motifs act with strong cooperativity. While wild-type Loc1p contains eight PUN motifs, a synthetic protein comprising ten copies of its most N-terminal PUN motif (Loc1p (1-20)_10_) yielded an annealing activity comparable to the wild-type protein (**Figure 4G,H**). This finding suggests that the iteration of PUN motifs is sufficient to recapitulate the folding-catalyst activity of wild-type Loc1p.

An important question is how the activity of PUN motifs as folding catalyst is achieved at the molecular level. Our NMR experiments showed that a single PUN motif interacts with dsRNA, whereas peptides without a motif show considerably weaker binding (**Figure 4A-D**, **Figure S4A**). The secondary structure of a dsRNA is not significantly altered upon PUN motif binding but the peptide itself gains rigidity (**Figure 4F**). Furthermore, the binding of the PUN motif results in the thermal stabilization of dsRNA (**Figure 4C,D**). Our ^31^P NMR experiments are consistent with a direct interaction of the positively charged arginine/lysine side chains of the PUN motif with the phosphate groups of the dsRNA (**Figure S4B**). The interaction still appears quite dynamic and no specific interaction is formed. This observation for PUN motifs is in contrast with other peptides that have been shown to bind to RNA hairpins (70,71). The dynamic and sequence-unspecific interaction is consistent with a potential chaperoning function of the PUN motif.

It has been shown that Loc1p and Puf6p form a ternary complex with the ribosomal protein Rpl43p and facilitate its loading onto the 60S ribosomal subunit (12,13). After this step, Loc1p and Puf6p are released. It is conceivable that Loc1p is required to assist correct folding of rRNAs to allow Rpl43p to be positioned correctly on the ribosomal subunit.

One interesting feature of PUN motifs is that each of the three repetitive units of 1-3 positively charged residues ([RK](1,3)) is interspaced by 1-3 residues that are not positively charged ({RK}(1,3)) (**Figure 3E**, **Figure S3D**). Such spacing of positively charged side chains in PUN motifs is consistent with the spatial requirements for their interaction with the phosphoribose backbone of RNA (**Figure 8**). For example, the C_α_ atoms of two lysine residues spaced by one additional amino acid (e.g. K-X-K; **Figure 8A**) in an extended conformation span a distance of 6-7 Å. In fact, this is also the distance range between two neighboring phosphates of the phosphoribose moiety in single-and double-stranded DNA/RNA. In case of two adjacent lysines or arginines (e.g. K-K), their flexible, positively charged side chain may also bind to two consecutive phosphate groups (cis configuration, **Figure 8B**) or even phosphate groups of two different DNA or RNA strands (trans configuration, **Figure 8C**). Finally, a sequence of at least three consecutive lysine or arginine side chains (e.g. K-K-K; **Figure 8D**) may well explain the observed chaperone-like activity of PUN motifs. Since side chains of neighboring amino acids in an extended conformation usually point in opposite directions, two RNA/DNA strands may be connected by an interjacent amino acid stretch. The repetitive nature of PUN motifs in certain proteins allows for multivalent interactions, which is also suitable to explain the observed RNA-dependent phase separation of these proteins.

**Figure 8:**
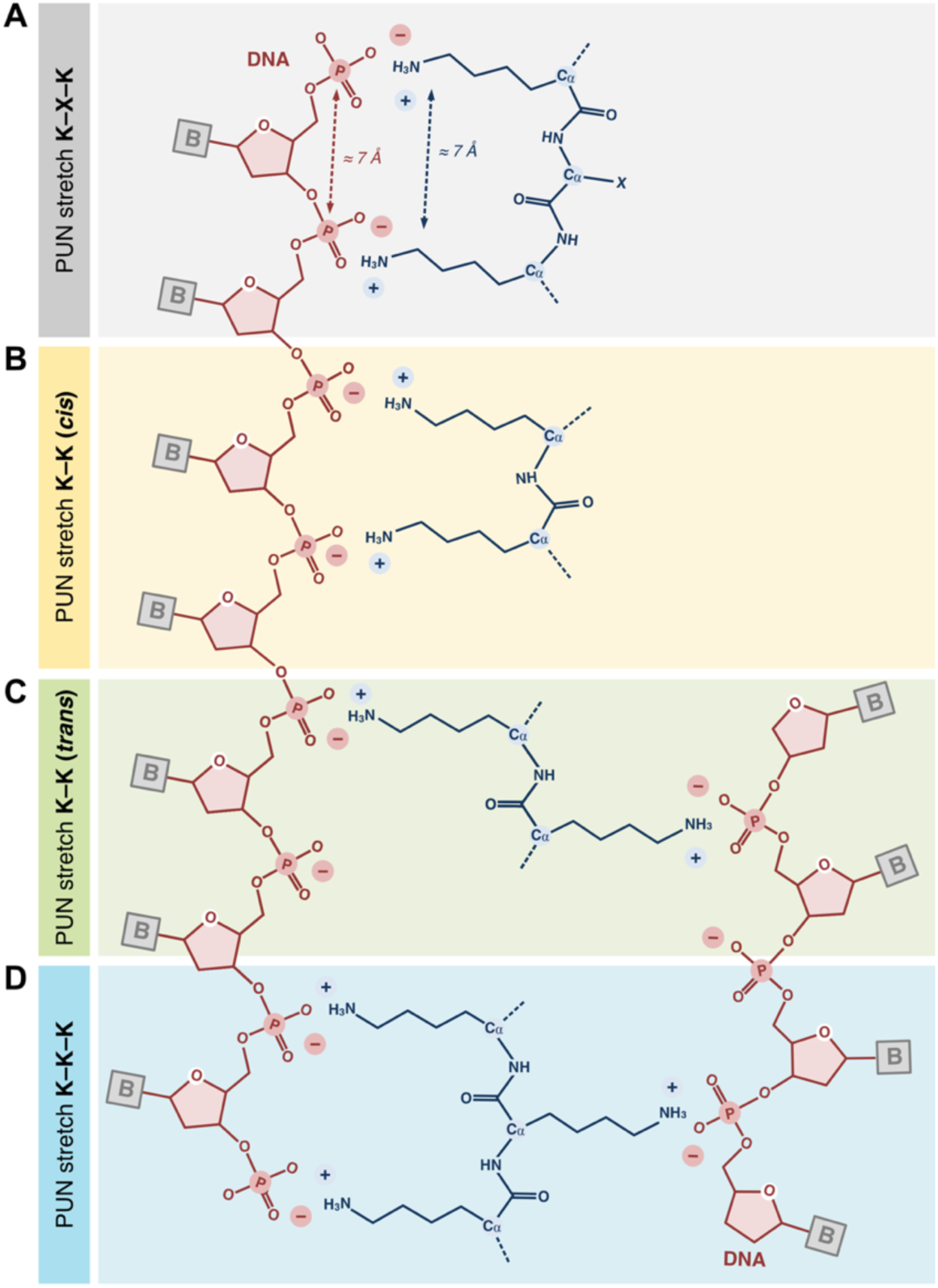
Schematic model of possible interaction modes of PUN motifs with nucleic acids. Sequence-unspecific interactions with phosphor-diester groups of nucleic-acids are mediated by positively charged side chains. **(A)** Two positively charged amino acids separated by a single non-positively charged amino acid have a distance that is very similar to the spacing of two neighboring phosphodiester groups in nucleic acids. Therefore, these two amino acids can potentially interact with two adjacent phosphor-diesters. **(B)** Due to the lengths and flexibilities of the side chains of arginine and lysine two neighboring amino acids can also interact with adjacent phosphodiesters of the same molecule (cis). **(C)** When pointing in opposite directions, two neighboring side chains can contact phosphodiester groups of two nucleic acid molecules (trans). **(D)** In a stretch of three consecutive arginines or lysines, these amino acids will also adopt a trans conformation. This feature should allow for the simultaneous contact of two nucleic-acid molecules by one PUN motif. This arrangement likely constitutes an important feature for higher organization principles, including RNA annealing and RNA-dependent phase transition.

This sequence-unspecific binding mode suggests that the motif fulfills a very general molecular function and therefore might be a ubiquitous motif for sequence-independent RNA interactions. The folding-catalyst activity and dsRNA-stabilizing functions of PUN motifs could in principle also support ribozymes other than the ribosome. Furthermore, the ability to undergo RNA-driven phase separation would potentially help such ribozymes to organize themselves in distinct functional foci in the cell, for instance, to locally concentrate their activities or recruit additional accessory factors.

In fact, we observed phase separation of Loc1p when mixed at concentrations of 5 µM or higher with total HeLa RNA. Since the formation of condensates depended on the protein concentration we wondered if the intra-nucleolar Loc1p concentration *in vivo* may be high enough to allow for phase separation. According to two different quantitative mass spectrometry studies, the theoretical nuclear Loc1p concentration is between 2 and 8 µM, calculated from the determined number of Loc1p molecules per cell and an average nuclear volume of 3 µm^3^ (72,73). Taking into account that the nucleolus occupies about one-third of the nuclear volume in yeast (74) and the predominant nucleolar localization of Loc1p, the concentration in the nucleolus is estimated to be between 6 and 24 µM. Considering the formation of condensates at 5 µM Loc1p in our *in vitro* assay, it seems reasonable to speculate that Loc1p can also form biomolecular condensates with RNA in the yeast nucleolus at the calculated concentrations.

It is conceivable that the RNA-dependent phase separation of Loc1p helps increasing its local concentration in the nucleolus to a level where it catalyzes correct folding of rRNA to allow for efficient joining of Rpl43p in the ribosomal maturation process. Such a scenario is consistent with the observation that in a *loc1Δ* strain reduced 60S subunit levels (4) and a slow-growth phenotype are observed (**Figure 1A**) (15). Furthermore, the observed Loc1p-dependent folding catalysis and phase separation of the E3 localization element of the *ASH1* mRNA (**Figure S6**) indicates that Loc1p might assist co-transcriptional folding of transported RNAs into their energetically most favored state. Of note, the RNA polymerase II-associated mediator complex has been shown to assemble into phase-separated co-transcriptional complexes (75). It is tempting to speculate that for folding catalysis of the nascent *ASH1* mRNA chain, the interaction of Loc1p with Spt5 (69) is supported by its phase-separating properties in the context of the RNA polymerase II-associated mediator complex.

In addition to Loc1p, other yeast proteins involved in RNA metabolism have been found to harbor RNA binding and annealing activities, which are not mediated by conventional/canonical, folded RNA-binding domains. For example, the mRNP packaging factor Yra1p and the mRNP biogenesis factor Tho2p possess RNA annealing activities and contain intrinsically disordered regions (IDRs) interspersed with clusters of positively charged amino acids. Interestingly, the RNA-annealing properties of Yra1p and Tho2p lie in (map to) these positively charged IDRs which contain three and four PUN motifs, respectively (76–79).

EMSAs showed that the PUN protein Loc1p does not discriminate between RNA and DNA binding (**Figure S1B,C**). It is therefore not surprising that proteins bearing PUN motifs are also found in DNA-related processes. For example, components of the chromatin remodeling ISWI complex, different DNA helicases and topoisomerases show enrichment of PUN motifs in *S. cerevisiae* as well as *D. melanogaster* and humans (**Tables S6**, **S8** and **S10**). This conclusion is further supported by the fact that *S. cerevisiae*, *D. melanogaster,* and even human histone H1 proteins contain multiple PUN motifs (**Table S10**). Their predominantly unstructured regions are thought to interact via basic residues with chromatin thus neutralizing the negative charge (80). Together with the globular domain of the histone 1, this charge neutralization allows for the condensation of chromatin (81). The observation that PUN motifs mediate nucleic acid-dependent phase separation suggests that transcriptional regulation also involves distinct DNA-protein phases as proposed recently (82–84).

Of note, histone 1 is subjected to posttranslational modifications like phosphorylation, acetylation and methylation (85,86). These modifications often occur within or in close proximity to PUN motifs, for instance in human histone H1.4 (85) or in *S. cerevisiae* histone H1 (87–89). A negative charge like a phospho-serine/-threonine or the removal of the positive charge of lysines by acetylation (acetyllysine) within or in close proximity to a PUN motif would alter the charge of this region and thereby might decrease its affinity to nucleic acids. We observed a similar affinity-modulating effect in PUN motifs of Loc1p. Here, negatively charged amino acids within or in close proximity to a PUN motif decreased its affinity to RNA (**Figure S2**). Thus, post-translational modification might offer a way to modulate nucleic acid binding affinities of proteins containing multiple PUN motifs. This model is supported by a recent study, showing that lysine-acetylation in disordered proteins can reverse RNA-dependent phase separation (90).

In the human proteome, we observed a considerable fraction of yeast PUN-proteins to be conserved (**Figure S9**). Furthermore, the strong accumulation of nucleic acid-related processes in the functional enrichment analysis of human PUN proteins (**Figure S8F**) indicates a functional conservation of paralogs between distant species. Our observation that up to 15% of the mRNA-interacting human proteins (91,92) contain three or more PUN motifs (**Table S10**), indicates that they play important roles in a large portion of the human RNA-binding proteins.

An important feature of PUN motifs is that they lack secondary structures. Thus, for PUN motifs to be active they would likely need to be located in disordered regions of these human proteins. Indeed, a systematic investigation of the secondary-structure environment shows that proteins containing five or more PUN motifs display a higher degree of disorderedness compared to other proteins, reflecting the preference of the motifs for unfolded regions (**Figure S8C**).

It has been proposed already several years ago that in an ancient RNA-dominated world short peptides might have helped to protect RNA against degradation and to stabilize defined RNA conformations (**Figure 9**) (93). This might have fostered the co-evolution of both types of molecules into functional RNPs. Accordingly, peptides are thought to have extended the structural and functional capabilities of RNAs and to facilitate the evolution of large RNPs like ribosomes (94,95). It is therefore easy to envision how in an ancient RNA-dominated world ribozymes might have benefited from short, stabilizing peptides, such as the PUN motif. Several years ago, Herschlag and colleagues provided experimental support for this hypothesis. A hammerhead ribozyme, whose catalytic activity was impeded by mutations, was reactivated to cleave its substrate when a positively charged peptide was added (96).

**Figure 9:**
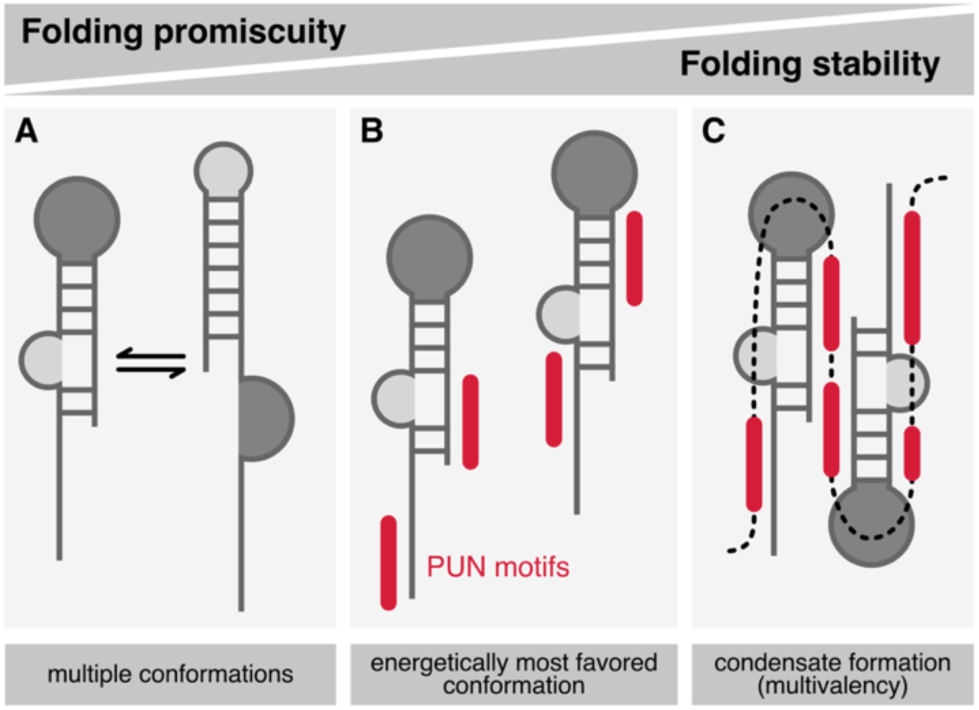
Model of the effect of isolated and repeating PUN motifs on promiscuous folding of RNA molecules. **(A)** Isolated RNA (schematic stem loop) may exist in an equilibrium of different secondary structures. **(B)** Binding of individual PUN motifs (red) to an RNA molecule stabilizes its energetically favored conformation. **(C)** A peptide chain with several PUN motifs acts cooperatively on RNA folding and facilitates the formation of condensates caused by their multivalent interactions.

However, it is much less clear what the direct benefit would have been in producing longer peptides that only later fold into small globular domains with more elaborate functions. For PUN motifs we observed that the fusion of two entities already induces strong cooperativity with greatly enhanced RNA-annealing activity. Using the PUN motifs as a paradigm, it is tempting to speculate that the potentiation of such basic functions by multimerization provided an immediate evolutionary benefit that favored the production of longer peptide chains and helped to pave the way into a world where functional RNPs might have replaced an RNA-dominated world. Our study offers the insight that such motifs still fulfill similar functions in today’s modern ribozymes, like ribosomes, spliceosomes or RNases. For example, proteins associated with the yeast ribosome, such as ribosomal proteins L15 and L24 or numerous rRNA processing factors CGR1, RRP14, RRP5, RRP8, RRP3 and the ribosomal biogenesis protein BMS1 contain several PUN motifs. Similarly, the pre-mRNA splicing factors 8, CLF1 and YJU2 as well as the RNaseP subunit POP1 include PUN motifs.

Furthermore, our results show that small RNA-binding motifs act cooperatively to catalyze RNA-based processes and at the same time can organize themselves into functional entities by phase separation (**Figure 9**). The latter has been described as organizing principle for a variety of cellular processes including the formation of nuclear and cytosolic bodies, such as paraspeckles, the nucleolus and stress granules (97–100). In an ancient world, repetitive elements such as PUN motifs would have added the benefit of organizing RNA-protein complexes into higher-order entities via phase separation and thus more efficient catalysis of RNA-based reactions. It will be interesting to find out how prevalent short, disordered RNA-binding motifs with dual functions, such as RNA annealing and phase separation, are and which essential cellular functions they carry out in today’s cells.

## Data availability

Uncropped images are available in the Supplementary Material. Microarray data is available for download via the PUMA database (http://puma.princeton.edu) experiment set No. 7342 and via Gene Expression Omnibus (https://www.ncbi.nlm.nih.gov/geo) experiment set No. GSE273374. Coordinates and structure factors have been deposited in the Protein Data Bank with the accession code 6YMC. The NMR chemical shift assignment of loc1p(3–15) has been deposited at the BMRB with accession code 52416. Supplementary Data are available at NAR Online.

## Funding

This work was supported by the Deutsche Forschungsgemeinschaft through grants to DN (DFG FOR855 and DFG FOR2333) and KZ (DFG FOR2333). DD was supported by the Emmy Noether Programme of the DFG (project number 246137224).

## Acknowledgments

We like to thank Vera Roman for her constant lab support and John Mattese from PUMAdb (http://puma.princeton.edu) for his important contribution. We acknowledge the use of the X-ray crystallography platform at the Institute of Structural Biology, Molecular Targets and Therapeutics Center, Helmholtz Munich. Moreover, we would like to thank the Core Facility Light Microscopy of the Medical Faculty at Ulm University and Dr. Christian Bökel for providing his expertise, and access to instrumentation funded by the Deutsche Forschungsgemeinschaft (DFG, German Research Foundation) – project number 257897648.

## Conflict of Interest Statement

The authors declare no competing interests.

## Notes

### Competing Interest Statement

The authors have declared no competing interest.

### Summary of Updates

We have added new results (Figure S6) and additional analyses (Figure S9).

