## Supplemental Figures and Tables for "Intrinsically disordered RNA-binding motifs cooperate to catalyze RNA folding and drive phase separation"

### Supplementary Information – Niedner-Boblenz et al.

**A** Micro-array analysis of RNAs immuno-precipitated with TAP-Loc1p

| Gene | Mean | Function |
|---|---|---|
| RDN58-1 | 3.803 | 5.8S rRNA |
| YLR161W | 3.707 | uncharacterized ORF in proximity to rDNA |
| RDN5-1 | 3.367 | 5S rRNA |
| YLR159W | 3.180 | uncharacterized ORF in proximity to rDNA |
| YKL084W | 2.080 | HOT13; mitochondrial intermembrane space protein |
| YHL041W | 2.057 | dubious ORF unlikely to encode a functional protein |
| YLR119W | 2.007 | SRN1; component of ESCRT-I; perhaps involved in RNA export |
| YCR018C | 1.977 | SRD1; involved in processing of pre-rRNA to mature rRNA |
| YEL021W | 1.957 | URA3; enzyme involved in biosynthesis of pyrimidines |
| RDN25-1 | 1.887 | 25S rRNA |

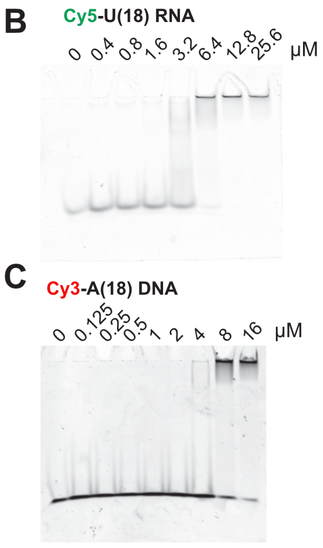


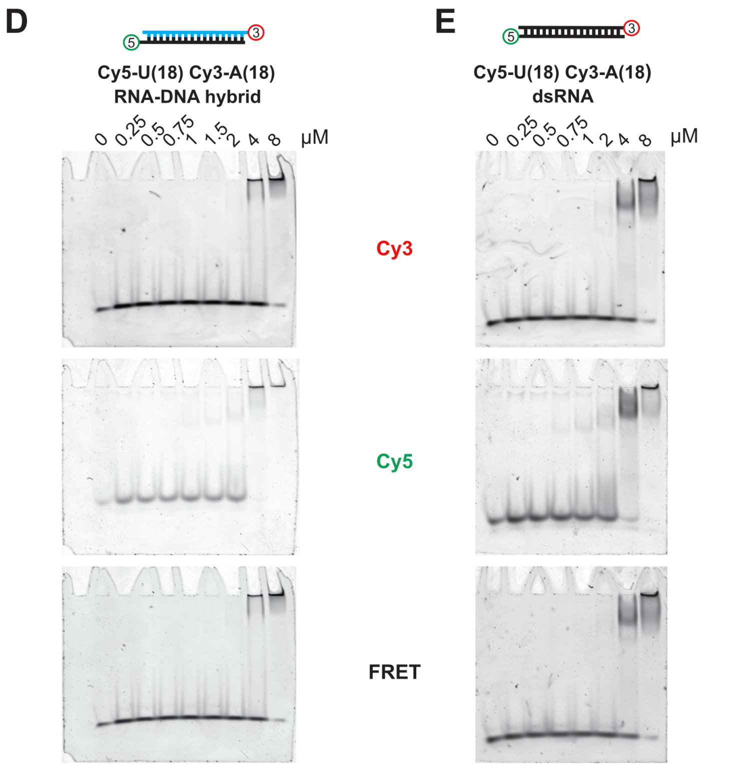


**Figure S1: Unspecific RNA binding by Loc1p. (A)** Microarray analysis of RNAs immunoprecipitated with TAP-tagged Loc1p revealed binding to rRNAs of the large ribosomal subunit. This table includes the ten most abundant features (including ORF and ncRNAs) that were detected in at least two out of three immunoprecipitations. Microarray data is available for download via the PUMA database (http://puma.princeton.edu), experiment set No. 7342. **(B-E)** Loc1p binds unspecifically to single-stranded and double-stranded nucleic acids. EMSAs with fluorescence-labeled nucleic acids show that Loc1p binds to single-stranded RNA (B) with a similar affinity as it binds to single-stranded DNA (C). In-gel FRET experiments (D,E) show that Loc1p binds to double-stranded RNA as well as to an RNA-DNA hybrid. For fluorescence EMSAs 100 nM of labeled RNA or DNA and the indicated amount of MBP-Loc1p were incubated at RT for 25 min before complexes were resolved on a 6 % TBE-PAGE. Fluorescence signals were detected using a typhoon scanner.

| No | Sequence | Binding | (R,K) / (R,K,D,E) | Motif |
|---|---|---|---|---|
| A1 | MAPKKPSKRQNLRREVAPEV | ++ | 75% | + |
| A2 | KKPSKRQNLRREVAPEVFQD | ++ | 67% | + |
| A3 | SKRQNLRREVAPEVFQDSQA | ++ | 57% | |
| A4 | QNLRREVAPEVFQDSQARNQ | | 50% | |
| A5 | RREVAPEVFQDSQARNQLAN | | 50% | |
| A6 | VAPEVFQDSQARNQLANVPH | | 33% | |
| A7 | EVFQDSQARNQLANVPHLTE | | 25% | |
| A8 | QDSQARNQLANVPHLTEKSA | | 50% | |
| A9 | QARNQLANVPHLTEKSAQRK | +++ | 80% | |
| A10 | NQLANVPHLTEKSAQRKPSK | +++ | 80% | + |
| A11 | ANVPHLTEKSAQRKPSKTKV | +++ | 83% | + |
| A12 | PHLTEKSAQRKPSKTKVKKE | +++ | 78% | + |
| A13 | TEKSAQRKPSKTKVKKEQSL | +++ | 78% | + |
| A14 | SAQRKPSKTKVKKEQSLARL | +++ | 86% | |
| A15 | RKPSKTKVKKEQSLARLYGA | +++ | 88% | |
| A16 | SKTKVKKEQSLARLYGAKKD | +++ | 78% | |
| A17 | KVKKEQSLARLYGAKKDKKG | +++ | 78% | |
| A18 | KEQSLARLYGAKKDKKGKYS | +++ | 78% | + |
| A19 | SLARLYGAKKDKKGKYSEKD | +++ | 70% | + |
| A20 | RLYGAKKDKKGKYSEKDLNI | ++ | 70% | + |
| B1 | GAKKDKKGKYSEKDLNIPTL | ++ | 67% | + |
| B2 | KDKKGKYSEKDLNIPTLNRA | +++ | 67% | + |
| B3 | KGKYSEKDLNIPTLNRAIVP | ++ | 67% | |
| B4 | YSEKDLNIPTLNRAIVPGVK | + | 75% | |
| B5 | KDLNIPTLNRAIVPGVKIRR | +++ | 83% | |
| B6 | NIPTLNRAIVPGVKIRRGKK | +++ | 100% | + |
| B7 | TLNRAIVPGVKIRRGKKGKK | +++ | 100% | + |
| B8 | RAIVPGVKIRRGKKGKKFIA | +++ | 100% | + |
| B9 | VPGVKIRRGKKGKKFIADND | +++ | 78% | + |
| B10 | VKIRRGKKGKKFIADNDTLT | +++ | 78% | + |
| B11 | RRGKKGKKFIADNDTLTLNR | +++ | 78% | + |
| B12 | KKGKKFIADNDTLTLNRLIT | +++ | 71% | |
| B13 | KKFIADNDTLTLNRLITTIG | | 75% | |
| B14 | IADNDTLTLNRLITTIGDKY | | 40% | |
| B15 | NDTLTLNRLITTIGDKYDDI | | 33% | |
| B16 | LTLNRLITTIGDKYDDIAES | | 33% | |
| B17 | NRLITTIGDKYDDIAESKLE | | 38% | |
| B18 | ITTIGDKYDDIAESKLEKAR | | 44% | |
| B19 | IGDKYDDIAESKLEKARRLE | | 45% | |
| B20 | KYDDIAESKLEKARRLEEIR | + | 50% | |
| C1 | DIAESKLEKARRLEEIRELK | | 50% | |
| C2 | ESKLEKARRLEEIRELKRKE | + | 57% | |
| C3 | LEKARRLEEIRELKRKEIER | + | 57% | + |
| C4 | ARRLEEIRELKRKEIERKEA | + | 57% | + |
| C5 | LEEIRELKRKEIERKEALKQ | + | 54% | + |
| C6 | IRELKRKEIERKEALKQDKL | ++ | 62% | + |
| C7 | LKRKEIERKEALKQDKLEEK | ++ | 57% | + |
| C8 | KEIERKEALKQDKLEEKKDE | | 47% | + |
| C9 | ERKEALKQDKLEEKKDEIKK | | 53% | + |
| C10 | EALKQDKLEEKKDEIKKKSS | + | 57% | + |
| C11 | KQDKLEEKKDEIKKKSSVAR | ++ | 62% | + |
| C12 | KLEEKKDEIKKKSSVARTIR | ++ | 67% | + |
| C13 | EKKDEIKKKSSVARTIRRKN | +++ | 75% | |
| C14 | DEIKKKSSVARTIRRKNKRD | ++ | 75% | + |
| C15 | KKKSSVARTIRRKNKRDMLK | +++ | 91% | + |
| C16 | SSVARTIRRKNKRDMLKSEA | +++ | 78% | + |
| C17 | ARTIRRKNKRDMLKSEAKAS | ++ | 80% | + |
| C18 | IRRKNKRDMLKSEAKASESK | +++ | 73% | + |
| C19 | KNKRDMLKSEAKASESKTEG | + | 60% | + |
| C20 | RDMLKSEAKASESKTEGRKV | + | 60 | |
| D1 | LKSEAKASESKTEGRKVKKV | +++ | 70% | + |
| D2 | EAKASESKTEGRKVKKVSFA | +++ | 67% | + |
| D3 | AKASESKTEGRKVKKVSFAQ | ++ | 75% | + |

| normalized intensity | Binding |
|---|---|
| 10 - 39 % | + |
| 40 - 69 % | ++ |
| 70 - 100 % | +++ |

**Figure S2**: **Peptides used in the peptide tiling array and quantified binding intensities.** Analysis of *ASH1* E3 binding to the Loc1p peptide tiling array. Shown are the position in the array, the peptide sequence, normalized binding intensity for each position, ratio of positively charged residues versus charged residues, and whether the respective peptide contains a repetitive, positively charged motif. The peptide with the highest intensity was set to 100 % and other peptides were normalized accordingly. Peptides with a higher portion of negatively charged residues show lower intensities and thus binding.

**A**

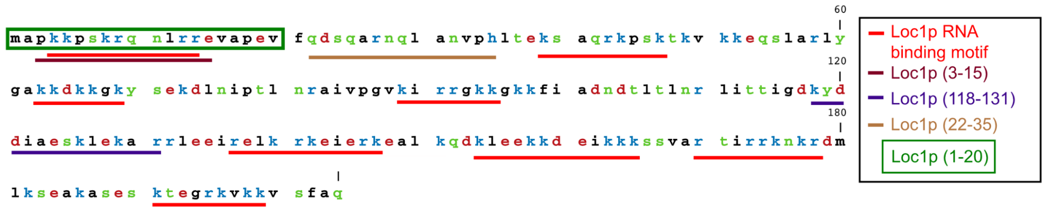


**B**

| Loc1p: biochemical data | |
|---|---|
| Amino acids | 204 |
| Molecular weight | 23.6 kDa |
| Isoelectric point | 10.3 |
| No of + charged residues | 61 |
| No of - charged residues | 30 |

**C**

| Name | Sequence | motif | Assay |
|---|---|---|---|
| Loc1p (3-15) | pkkpskrqnlrre | 1 | NMR |
| Loc1p (118-131) | kyddiaesklekar | 0 | NMR |
| Loc1p (22-35) | qdsqarnqlanvph | 0 | NMR |
| Loc1p (1-20) | mapkkpskrqnlrrevapev | 1 | Chaperone assay |
| Loc1p $(1\text{-}20)_2$ | mapkkpskrqnlrrevapevmapkkpskrqnlrrevapev | 2 | Chaperone assay |

**D**

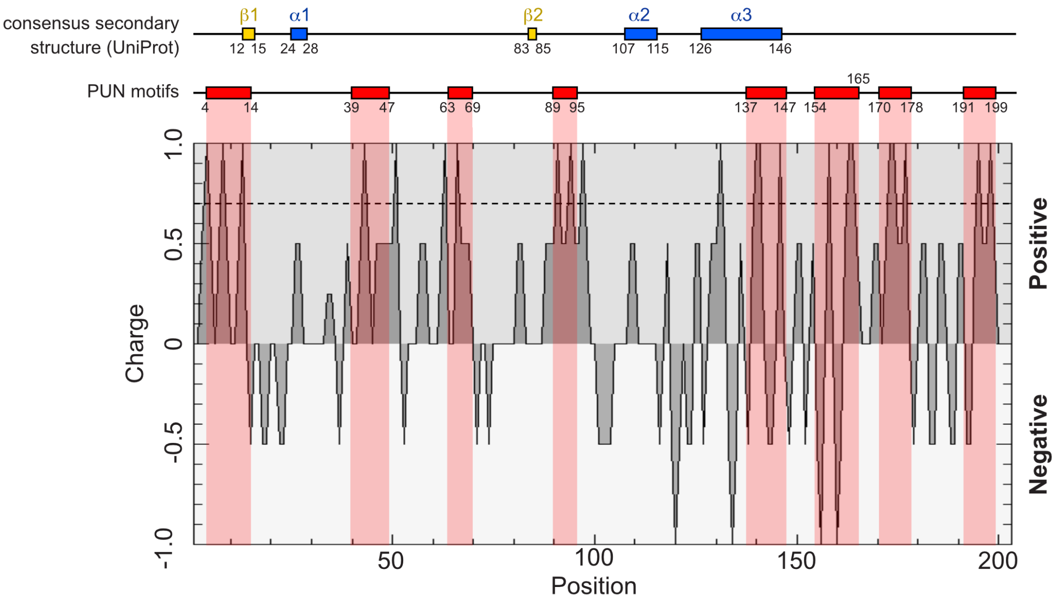


**Figure S3**: Protein regions of Loc1p interacting with RNA and PUN motifs. **(A)** Amino acid sequence of Loc1p colored according to their electrochemical properties (charges: negative in red, positive in blue, polar neutral in green, polar hydrophobic in black). RNA binding motifs within the Loc1p sequence are underlined in red. **(B)** Loc1p protein parameters derived from www.expasy.org/proteomics. **(C)** List Loc1p-derived peptides used in this study. The RNA binding motif is depicted in red and the experiment is indicated. The sequences of the used peptides are also indicated in (B). **(D)** Charge plot of the Loc1p sequence. The charge is plotted against the amino acid sequence of Loc1p (window size: 2 amino acids). Above the charge plot, schematic representations of the Loc1p sequence illustrate the location of secondary structure elements (as described in UniProt) and PUN repeats that match the consensus sequence. Note that each PUN repeat is characterized by positively charged amino acids while the spacers typically lack positive charges and instead bear one or more negatively charged residues.

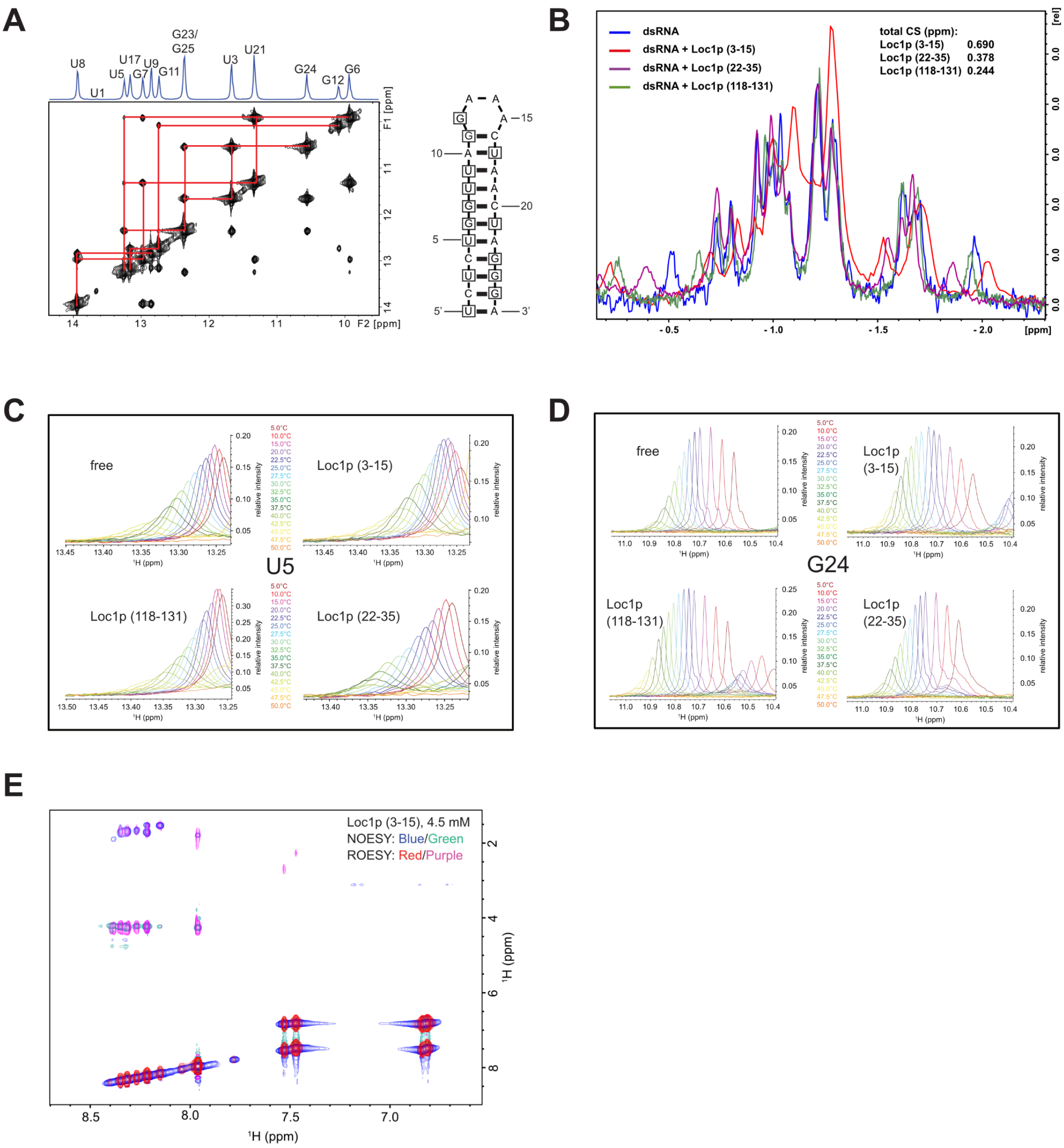


**Figure S4: NMR studies on the interaction between Loc1p and RNA. (A)** Complete imino signal assignment of the dsRNA. To the right the stem-loop is shown with assigned bases marked with a square. **(B)** Titration of all three peptides in 1D-$^{31}$P-NMR experiments with stem loop RNA. Although each peptide resulted in resonance shifts at high concentrations, they showed different effects. Resonances of phosphodiester bonds located in an A-type or B-type nucleic acid helix usually cluster around chemical shifts of -1 ppm, which can also be observed here (Gorenstein, 1984; Wüthrich, 1986). Terminal phoshomonoesters are shifted downfield and phosphodiesters of bases located at a terminal stem close to a loop are shifted upfield. The resonances (at -0.5 and -2 ppm) corresponding to terminally located phosphorus atoms exhibit the strongest shifts upon addition of peptides (5:1 ratio, peptide:RNA). Interestingly, Loc1p (22-35) and Loc1p (118-131) do not induce shifts on resonances clustered around -1 ppm, but Loc1p (3-15) does, indicating that Loc1p (3-15), the only one with chaperone activity and showing to stabilize the RNA strongest also binds to the backbone of nucleotides located at the core of the helix. **(C, D)** Selected, overlayed 1D imino NMR experiments of the RNA tetraloop free and bound to different peptides during temperature scans. (C) Zooms into the spectral region of the U5 imino peak of spectra of free RNA (upper left), bound to Loc1p(3-15) (upper right), Loc1p(118-131) (lower left), and Loc1p(22-35) (lower right). The peak positions have been altered manually to allow for a better comparison between signal intensities. The peak intensities for each temperature have been used to generate the melting curves and to estimate the melting temperature for U5 base pairing. Loc1p(3-15) stabilizes this base pair compared to free RNA, as the peak is still at its maxima at 25°C compared to 15°C in absence of peptide. The other two peptides on the contrary destabilize this base pair and signal intensity already decreases after raising the temperature above 10°C. (D) shows the same as (C) but for the imino peak of G24. This peak was more isolated and it was not necessary to adjust the peak position manually. Thus, their chemical shifts are the experimental ones. The color and corresponding temperatures are shown in between each spectrum. **(E)** Comparison of 2D ROESY and NOESY NMR spectra of Loc1p(3-15). In order to assess whether the lack of NOEs for Loc1p(3-15) in absence of RNA is due to lack of structure or due to its size and thus being at the zero-crossing for NOESY-based experiments, we compared a ROESY (no zero-crossing) and NOESY spectra. Even at 4.5 mM concentration, also the ROESY experiment shows only a few intra-residual NOE cross peaks, clearly demonstrating that Loc1p(3-15) is unstructured.

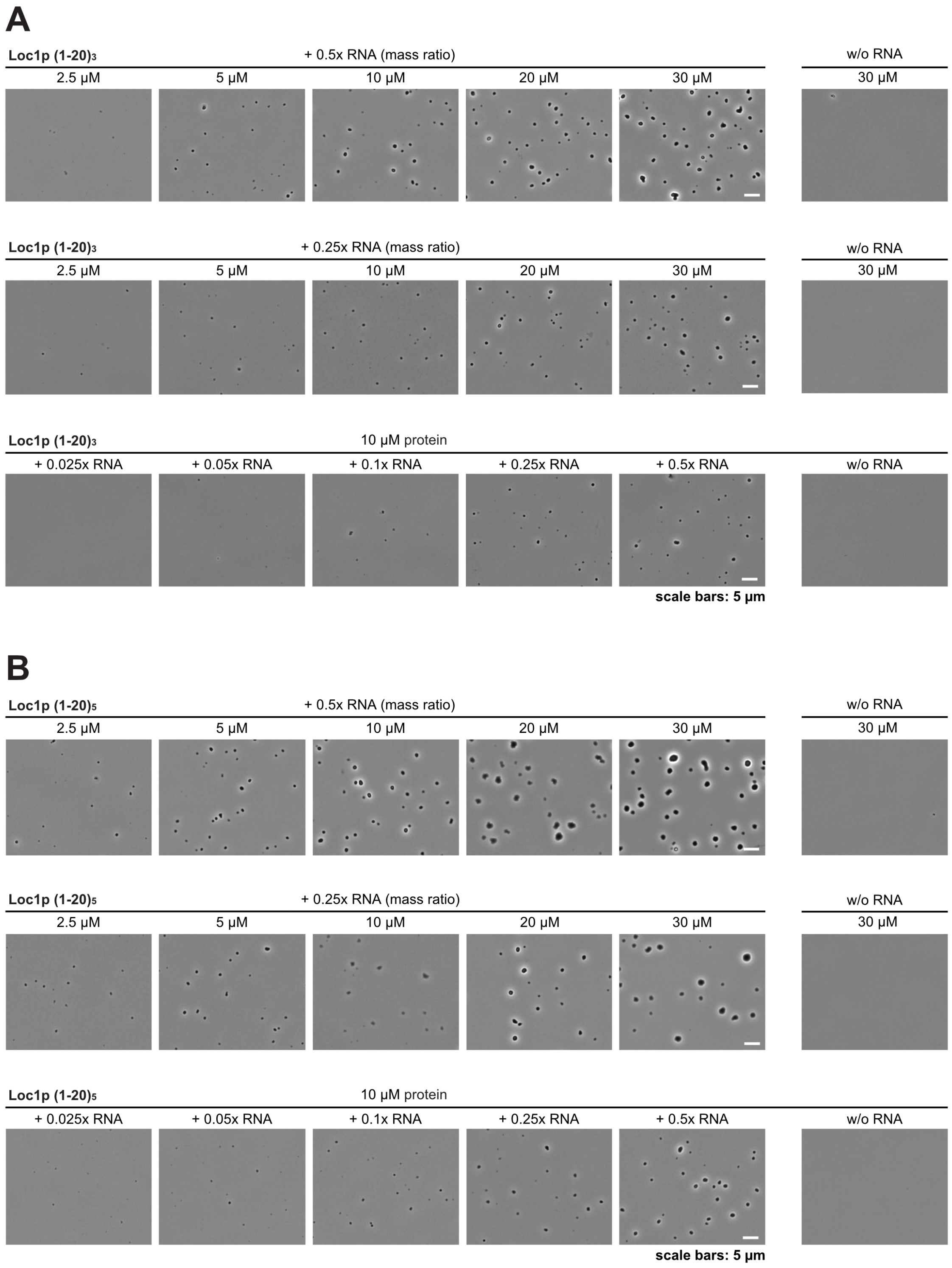


**Figure S5: Assessment of condensate formation by different PUN-motif containing proteins. (A-B)** RNA-dependent condensate formation by protein fragments containing three (A) or five (B) copies of the most N-terminal PUN motif of Loc1p. Condensate formation occurred already at lower concentrations when Loc1p (1-20)$_5$ was used instead of Loc1p (1-20)$_3$. No phase separation was observed in absence of RNA. The formation of condensates was assessed by phase-contrast microscopy.

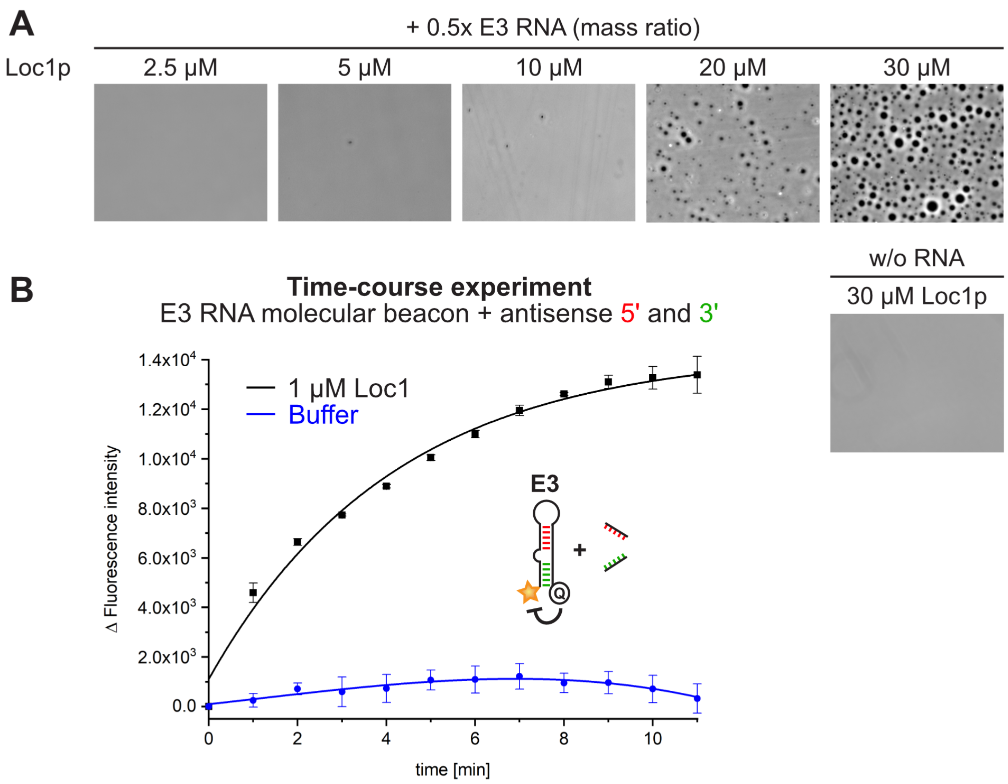


**Figure S6: Loc1p forms condensates with a 28nt E3 localization element (LE) of *ASH1* mRNA and promotes conformational changes in its stem-loop. (A)** Phase separation of full-length Loc1p is observed at concentrations of 5 µM or higher when mixed with the 28 nt E3 LE stem-loop, while no phase separation is observed without RNA. **(B)** Loc1p promotes conformational changes in the stem-loop of a 28nt E3 LE molecular beacon (5'-FAM and 3'-BMNQ-535). In the presence of two 5nt antisense RNAs (5'anti-E3 and 3'anti-E3), titration of Loc1p changes the fluorescence intensity of an E3 LE molecular beacon in time-course experiments, suggesting that Loc1p catalyzes the formation of the energetically most favored state.

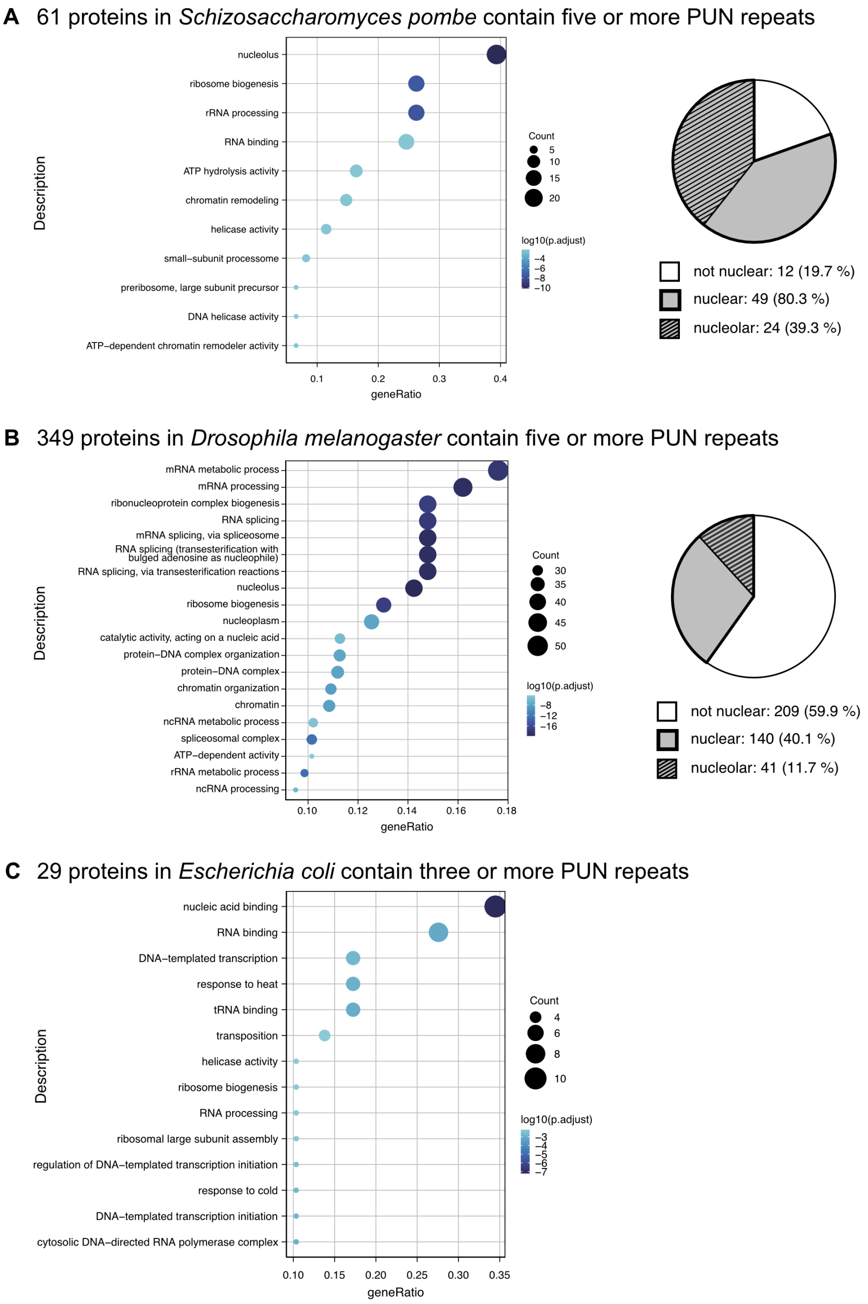


**Figure S7: Enrichment of PUN motif-containing proteins in distant eukaryotic and procaryotic species. (A)** GO-term analysis of 61 *Schizosaccharomyces pombe* proteins containing at least five PUN motifs (three of them in 150 aa) shows that pathways of ribosome biogenesis, RNA processing, and RNA binding are enriched. Analysis of the cellular localization of the 61 hits shows a clear preference for the nuclear and nucleolar compartments of the cell. **(B)** 349 *Drosophila melanogaster* proteins containing five or more PUN motifs with three motifs in at least 150 aa are also enriched in the nucleus. GO-term analysis of the cellular localization of the 349 hits shows a preference for the nuclear and nucleolar compartments of the cell. **(C)** We identified 29 *Escherichia coli* proteins containing at least 3 PUN motifs, which are involved in RNA metabolic processes. GO-term analysis of annotated processes shows that pathways of nucleic acid binding and transcription are enriched.

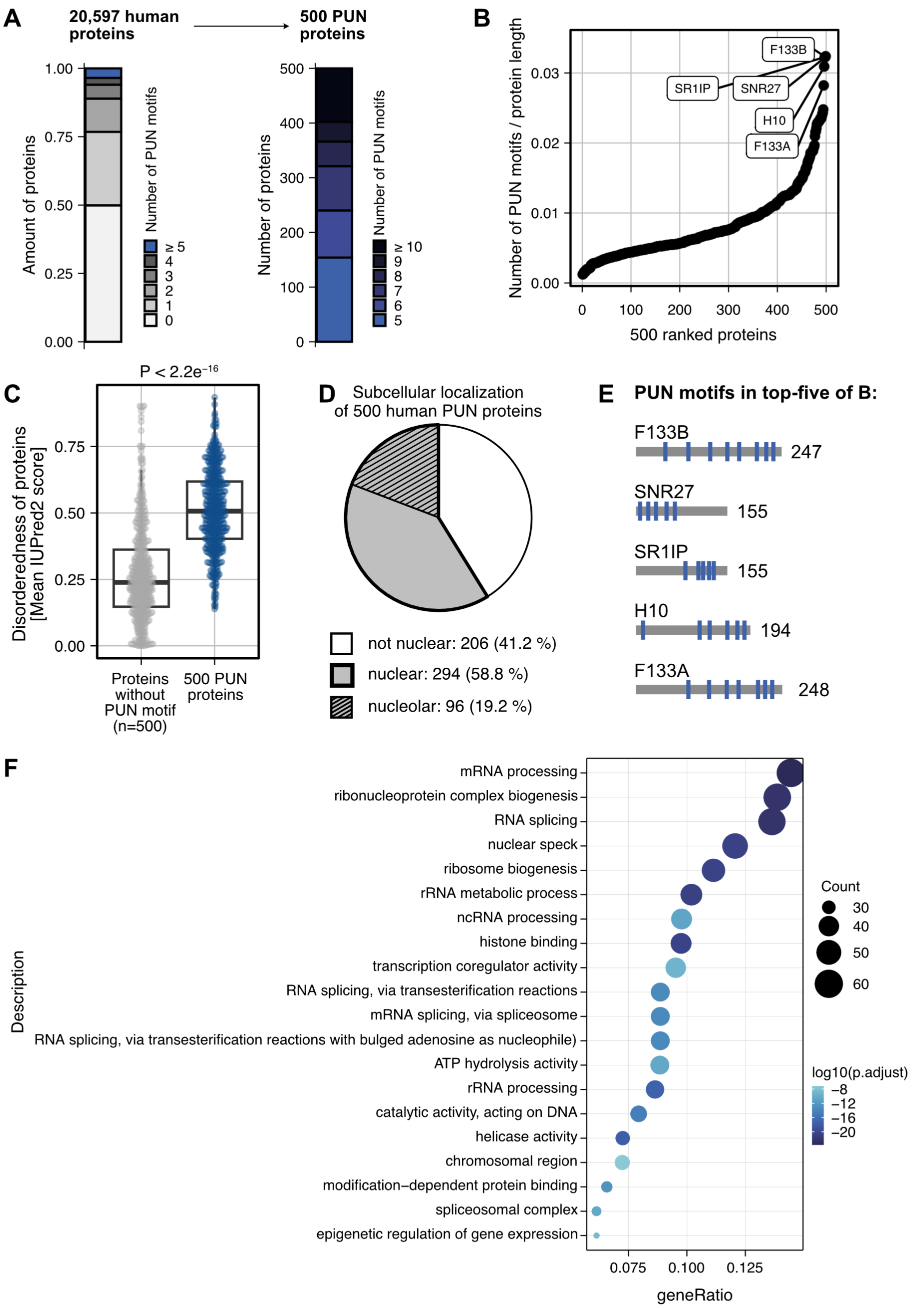


**Figure S8: Human PUN motif-containing proteins are enriched in the nucleus and involved in RNA-related processes. (A)** Identification of 500 human proteins that contain at least five PUN motifs in total and three in 150 amino acids. **(B)** PUN motif density in the 500 human PUN proteins. Proteins with the highest PUN motif density are involved in RNA-binding, DNA condensation, or splicing. The sperm head cytoskeletal calyx protein CYLC2 was removed because of its involvement in an RNA-unrelated process. **(C)** Human PUN proteins are characterized by a higher degree of disorderedness compared to proteins that do not match the criteria, reflecting the preference of PUN motifs for disordered regions. **(D)** Analysis of the intracellular distribution of the 500 hits shows a clear preference for the nuclear compartment of the cell (59 %). **(E)** Distribution of PUN motifs in top-hits from (B). **(F)** GO-term analysis of annotated processes ordered by P-values. The RNA processing pathways are enriched in human proteins bearing PUN motifs.

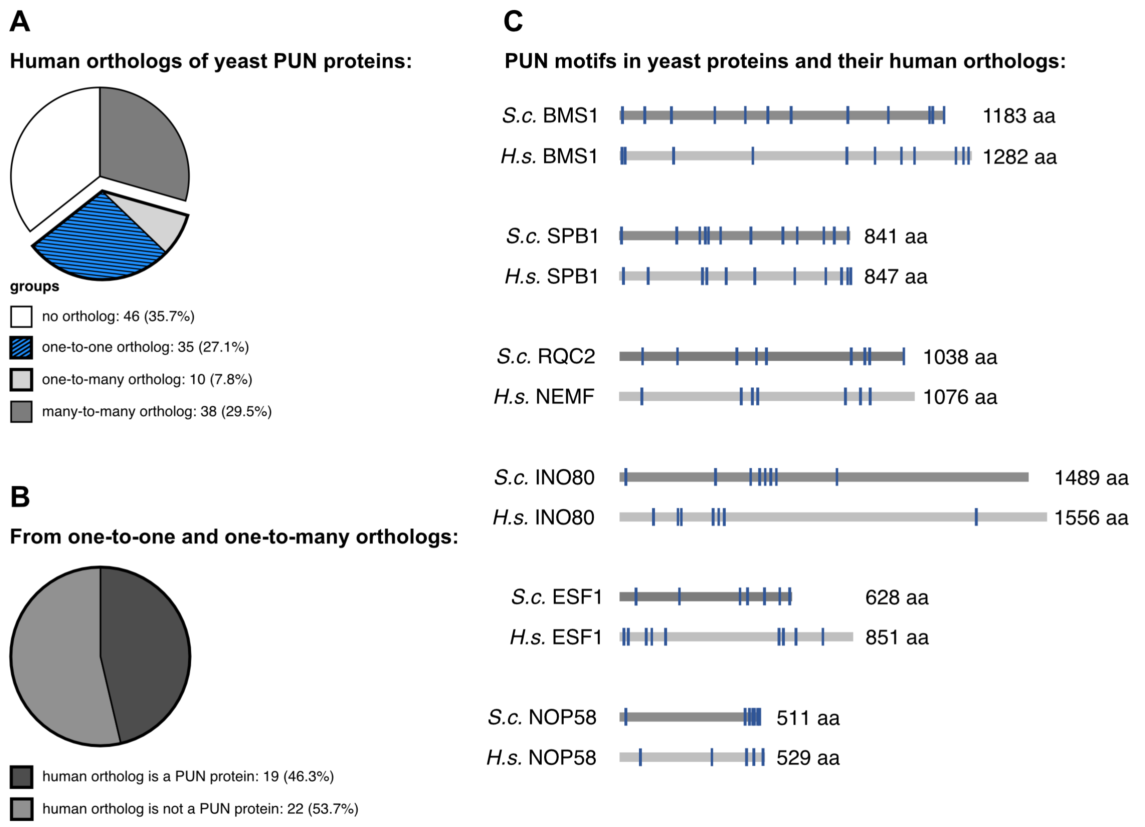


**Figure S9: Characterization of human orthologs of the 95 *S. cerevisiae* PUN proteins. (A)** A database search for human orthologs using the 95 yeast PUN proteins identified 35 one-to-one and 10 one-to-many orthologs, while 46 proteins have no human counterpart. **(B)** 46% of the one-to-one and one-to-many human orthologs are also PUN proteins in human, while 53% do not fulfill both criteria (5 or more PUN motifs and at least 3 PUN motifs within 150 amino acids). **(C)** Examples from the top-20 of yeast PUN proteins with their human orthologs. Protein length (in amino acids) is shown to scale in bar representation and the relative locations of PUN motifs are indicated by blue rectangles. Note that orthologous proteins are similar in length and clustering of the PUN motifs.

**Table S1**: **Isoelectric points of ribosomal proteins**, Theoretical isoelectric point calculated with ProtParam (www.expasy.org). Proteins with pI ≥ 9.0 are shown in green, between 8.9 and 7.0 in yellow (none) and with pI ≤ 6.9 in red.

| Large subunit | Size (aa) | pI |
|---|---|---|
| RPL1A/B | 217 | 10.51 |
| RPL2A/B | 254 | 11.74 |
| RPL3 | 387 | 11.1 |
| RPL4A/B | 362 | 11.4 |
| RPL5 | 297 | 6.82 |
| RPL6A | 176 | 10.91 |
| RPL6B | 176 | 10.89 |
| RPL7A/B | 244 | 10.97 |
| RPL8A | 256 | 10.85 |
| RPL8B | 256 | 10.83 |
| RPL9A | 191 | 10.54 |
| RPL9B | 191 | 10.47 |
| RPL10 | 221 | 10.84 |
| RPL11A/B | 174 | 10.73 |
| RPL12A/B | 165 | 10.2 |
| RPL13A | 199 | 11.79 |
| RPL13B | 199 | 11.71 |
| RPL14A/B | 138 | 11.63 |
| RPL15 | 204 | 11.96 |
| RPL16A | 199 | 11.28 |
| RPL16B | 198 | 11.34 |
| RPL17A/B | 184 | 11.62 |
| RPL18A/B | 186 | 12.24 |
| RPL19A/B | 189 | 11.93 |
| RPL20A/B | 172 | 11.12 |
| RPL21A/B | 160 | 11.2 |
| RPL22A | 121 | 6.01 |
| RPL22B | 122 | 5.98 |
| RPL23A/B | 137 | 11.11 |
| RPL24A | 155 | 11.9 |
| RPL24B | 155 | 11.97 |
| RPL25 | 142 | 10.94 |
| RPL26A | 127 | 11.35 |
| RPL26B | 127 | 11.25 |
| RPL27A/B | 136 | 11.16 |
| RPL28 | 149 | 11.4 |
| RPL29 | 59 | 11.99 |
| RPL30 | 105 | 10.64 |
| RPL31A/B | 113 | 10.8 |
| RPL32 | 130 | 11.83 |
| RPL33A/B | 107 | 11.76 |
| RPL34A/B | 121 | 11.56 |
| RPL35A/B | 120 | 11.36 |
| RPL36A/B | 100 | 12.15 |
| RPL37A | 88 | 12.19 |
| RPL37B | 88 | 12.32 |
| RPL38 | 78 | 11.65 |
| RPL39 | 51 | 12.55 |
| RPL40A/B | 128 | 10.63 |
| RPL41A/B | 25 | 12.96 |
| RPL42A/B | 106 | 11.38 |
| RPL43A/B | 92 | 11.21 |

| Small subunit | Size (aa) | pI |
|---|---|---|
| RPS0A | 252 | 4.47 |
| RPS0B | 252 | 4.51 |
| RPS1A | 255 | 10.79 |
| RPS1B | 255 | 10.81 |
| RPS2 | 254 | 11.21 |
| RPS3 | 240 | 10.22 |
| RPS4A/B | 261 | 10.9 |
| RPS5 | 225 | 9.11 |
| RPS6A/B | 236 | 11.22 |
| RPS7A | 190 | 10.64 |
| RPS7B | 190 | 10.73 |
| RPS8A/B | 200 | 11.42 |
| RPS9A | 197 | 10.8 |
| RPS9B | 195 | 10.9 |
| RPS10A | 105 | 9.63 |
| RPS10B | 105 | 9.92 |
| RPS11A/B | 156 | 11.55 |
| RPS12 | 143 | 4.5 |
| RPS13 | 151 | 11.24 |
| RPS14A | 137 | 11.43 |
| RPS14B | 138 | 11.28 |
| RPS15 | 142 | 11.45 |
| RPS16A/B | 143 | 11.08 |
| RPS17A/B | 136 | 11.28 |
| RPS18A/B | 146 | 11.06 |
| RPS19A/B | 144 | 10.45 |
| RPS20 | 121 | 10.33 |
| RPS21A/B | 87 | 5.9 |
| RPS22A/B | 130 | 10.76 |
| RPS23A/B | 145 | 11.47 |
| RPS24A/B | 135 | 11.29 |
| RPS25A/B | 108 | 11.13 |
| RPS26A | 119 | 11.49 |
| RPS26B | 119 | 10.6 |
| RPS27A/B | 82 | 9.51 |
| RPS28A/B | 67 | 11.42 |
| RPS29A | 56 | 11.06 |
| RPS29B | 56 | 10.8 |
| RPS30A/B | 63 | 12.24 |
| RPS31 | 152 | 10.67 |

| Stalk | Size (AA) | pI |
|---|---|---|
| RPP0 | 312 | 4.56 |
| RPP1A | 106 | 3.61 |
| RPP1B | 106 | 3.68 |
| RPP2A | 106 | 3.77 |
| RPP2B | 110 | 3.89 |
| RKM2 | 479 | 5.1 |

**Table S2**: **Microarray data showing Loc1p-bound RNAs.** List of most enriched RNAs when comparing mean of three Loc1p experiments with corresponding control experiments. YORF descriptions starting with "SPL2" refer to probes against intronic regions of a given gene (named thereafter), descriptions starting with "I" refer to probes against intergenic regions. The full set of dataset is available for download via the PUMA database (http://puma.princeton.edu; experiment #7342).

| YORF | Description | Loc1p #1 | Loc1p #2 | Loc1p #3 | Mean Loc1p | Rank | Mean wt |
|---|---|---|---|---|---|---|---|
| SPL2YML085C | TUB1; alpha-tubulin | 3,88 | | | 3,880 | 1,000 | 0,745 |
| RDN58-1 | 5.8S rRNA | 5,41 | 2,13 | 3,87 | 3,803 | 0,999 | 0,380 |
| IYLR161W | encoded near the ribosomal DNA region | 5,57 | 1,64 | 3,91 | 3,707 | 0,999 | 3,233 |
| RDN5-1 | 5S rRNA | 5,39 | 1,21 | 3,5 | 3,367 | 0,999 | 2,565 |
| IYLR159W | encoded near the ribosomal DNA region | 5,01 | 0,96 | 3,57 | 3,180 | 0,999 | 3,575 |
| SPL2YML025C | YML6; mitochondrial ribosomal protein | 2,92 | | | 2,920 | 0,999 | 0,525 |
| SPL2YLR406C | RPL31B; large ribosomal subunit protein | 2,66 | | | 2,660 | 0,999 | 0,545 |
| SPL2YJL225C | uncharacterized ORF | 2,65 | | | 2,650 | 0,998 | 0,630 |
| YNL204C | protein of unknown function | | | 2,5 | 2,500 | 0,998 | 0,110 |
| SPL2YIL052C | RPL34B; large ribosomal subunit protein | 3,64 | | 0,75 | 2,195 | 0,998 | 0,678 |
| YDR403W | Sporulation-specific enzyme | | 1,18 | 3,03 | 2,105 | 0,998 | 0,028 |
| YKL084W | HOT13; mitochondrial protein | 1,75 | 0,94 | 3,55 | 2,080 | 0,998 | -0,300 |
| YHL041W | dubious ORF | 1,59 | 2,19 | 2,39 | 2,057 | 0,998 | -0,158 |
| YGL183C | MND1; required for recombination | | 2,01 | | 2,010 | 0,997 | 0,633 |
| YOR343C | dubious ORF | | | 2,01 | 2,010 | 0,997 | 0,060 |
| YLR119W | SRN2; ESCRT-I component | 0,93 | 1,92 | 3,17 | 2,007 | 0,997 | -0,435 |
| SPL2YIL004C | BET1; v-SNARE | 3,08 | | 0,93 | 2,005 | 0,997 | 0,530 |
| YJL137C | GLG2; Glycogenin glucosyltransferase | | 1,6 | 2,36 | 1,980 | 0,997 | -0,015 |
| SPL2YMR225C | MRPL44; mitochondrial ribosomal protein | 2,98 | | 0,98 | 1,980 | 0,997 | 0,403 |
| YCR018C | SRD1; processing of pre-rRNA | 2,93 | 1,62 | 1,38 | 1,977 | 0,997 | -0,060 |
| YEL021W | URA3; OMP decarboxylase | 2,1 | 1,39 | 2,38 | 1,957 | 0,997 | 0,208 |
| YPR096C | protein of unknown function | | 0,55 | 3,34 | 1,945 | 0,996 | 0,070 |
| YJL103C | protein of unknown function | | 1,89 | | 1,890 | 0,996 | 0,293 |
| RDN25-1 | 25S rRNA | 2,62 | 1,4 | 1,64 | 1,887 | 0,996 | -1,288 |
| IYLRCDELTA8 | Ty1 long terminal repeat | 3,31 | -0,38 | 2,71 | 1,880 | 0,996 | 3,253 |
| YOR391C | HSP33; chaperone | | 1,18 | 2,56 | 1,870 | 0,996 | 0,088 |
| SPL2YLR078C | BOS1; v-SNARE | 1,54 | 1,05 | 3,02 | 1,870 | 0,996 | 0,565 |
| YJL135W | dubious ORF | | 1,78 | 1,89 | 1,835 | 0,995 | 0,070 |
| YGR217W | CCH1; calcium channel | 2,9 | 0,81 | 1,79 | 1,833 | 0,995 | -0,240 |
| YJL199C | dubious ORF | 1,64 | 1,91 | 1,91 | 1,820 | 0,995 | 0,000 |

**Table S5**: **Data collection and refinement statistics for the RNA hairpin tetraloop crystal structure**. Values in parentheses are for the highest-resolution shell.

| | |
|---|---|
| **Data collection** | |
| Beamline | SLS PXIII X06DA |
| Wavelength (Å) | 1.0000 |
| Space group | $P2_12_12_1$ |
| Cell dimensions *a*, *b*, *c* (Å) | 24.27, 52.43, 107.18 |
| Resolution (Å) | 50 – 2.0 (2.05 – 2.00) |
| $R_{merge}$ (%) | 9.2 (56.4) |
| $I / \sigma I$ | 13.1 (3.49) |
| CC (1/2) | 0.987 (0.681) |
| Completeness (%) | 97.0 (96.0) |
| Redundancy | 5.9 (5.7) |
| **Refinement** | |
| Resolution (Å) | 2.0 |
| No. reflections | 9,048 |
| $R_{work}$ / $R_{free}$ (%) | 17.5 / 24.1 |
| No. atoms | |
| RNA | 1,108 |
| Ba | 10 |
| Water | 97 |
| *B*-factor overall (Å$^2$) | 31.5 |
| R.m.s. deviations | |
| Bond lengths (Å) | 0.013 |
| Bond angles (°) | 2.28 |
| PDB ID | 6YMC |

#### External Tables for download:

**Table S3:** Microarray data showing all Loc1p-bound RNAs.

**Table S4:** List of nucleic acids used in this study.

**Table S6:** PUN motif proteins in *S. cerevisiae*.

**Table S7:** PUN motif proteins in *S. pombe*.

**Table S8:** PUN motif proteins in *D. melanogaster*.

**Table S9:** PUN motif proteins in *E. coli*.

**Table S10:** PUN motif proteins in *H. sapiens*.

**Table S11:** *S. cerevisiae* PUN motif proteins with their *H. sapiens* orthologs.

**Movie S1:** Time recording of Loc1p/RNA LLPS fusion.

**Movie S2:** Time recording of photobleaching and fluorescence recovery in a Loc1p/RNA LLPS FRAP experiment.
